# Integrated 5-HT_2A_–TrkB and G protein signaling in serotonergic psychedelic responses

**DOI:** 10.64898/2026.03.19.712961

**Authors:** Marco Taddei-Tardón, Lidia Medina-Rodríguez, Jessica L. Maltman, Sarah Hudson, Sritanvi Potukanuma, Javier Hidalgo Jiménez, Sandra M. Martín-Guerrero, Javier González-Maeso, Juan F. López-Giménez

**Affiliations:** lnstituto de Parasitologia y Biomedicina “Lopez-Neyra” (lPBLN-CSIC). E-18016 Granada. Spain; Department of Pharmacology and Toxicology, Virginia Commonwealth University School of Medicine, Richmond, VA 23298; Biomedical Sciences Graduate Program, Virginia Commonwealth University School of Medicine, Richmond, VA 23298

## Abstract

Serotonergic psychedelics have attracted considerable interest as promising therapeutic agents. However, the molecular mechanisms linking their acute hallucinogenic-like effects to longer-lasting neuroplastic responses remain incompletely understood, partly because of the scarcity of native neural models suitable for mechanistic studies. Here, we developed a neural stem cell-derived in vitro model capable of differentiating into neuronal and glial lineages and, after characterization, used it to investigate the molecular pharmacology of serotonergic psychedelics. A panel comprising tryptamines, phenethylamines and ergolines, including psychedelic compounds and selected non-psychedelic analogues, was evaluated alongside ketamine and TrkB agonists. Endpoints included dendritogenesis, synaptogenesis, immediate-early gene induction, BDNF expression and lactate production. TrkB silencing abolished dendritogenic responses to serotonergic psychedelics, ketamine and TrkB agonists, whereas 5-HT_2A_ receptor silencing selectively impaired serotonergic psychedelic-induced plasticity and altered TrkB-dependent responses. Most serotonergic compounds also increased synaptogenesis and induced *c-Fos* and *Egr-2* expression, although ligand-specific differences were evident, particularly for psilocin and the phenethylamines DOI and Ariadne. Uncoupling of G_q/11_ or G_i/o_ protein-dependent signaling differentially modified neuroplastic and transcriptional responses, indicating a ligand and endpoint dependent contribution of both pathways. Serotonergic psychedelics further induced a 5-HT_2A_ receptor dependent lactate response that was generally sensitive to disruption of either G_q/11_ or G_i/o_ protein coupling. Taken together, these findings support a model in which serotonergic psychedelics recruit an integrated 5-HT_2A_-TrkB signaling network with distinct structural, transcriptional and metabolic outputs, and establish this neural stem cell-derived system as a valuable platform for screening and dissecting the signaling basis of psychedelic action.

## Introduction

Psychedelic compounds have emerged in recent years as promising therapeutic tools for the treatment of various psychiatric conditions, particularly in cases where conventional interventions have proven ineffective, as exemplified by major depressive disorder[1]. Their therapeutic potential is largely attributed to the rapid onset of clinical response following administration, as well as to the persistence of therapeutic effects after a reduced number of initial doses [2, 3]. One of the most widely accepted hypotheses regarding their mechanism of action proposes that these compounds facilitate neuronal plasticity processes capable of reversing or restoring the structural atrophy observed in pyramidal neurons of the prefrontal cortex [4][5][6] as documented in post-mortem studies of patients affected by certain psychiatric pathologies[7]. This induction of neuronal plasticity, encompassing both morphological and functional aspects, is thought to be initiated by the binding of psychedelic compounds to membrane receptors, thereby activating multiple intracellular signaling cascades[8]. Beyond their effects on neuronal plasticity, psychedelic compounds are characterized by their capacity to induce transient and acute altered states of consciousness in human subjects[9]. This response has a well-established experimental correlate in preclinical animal models, most notably the head-twitch response assay (HTR), which is widely used due to the strong association reported between hallucinogenic effects in humans and this stereotyped behavioural response in rodents[10]. In the case of classical or serotonergic psychedelics, evidence suggests that their hallucinogenic, or more broadly psychoactive, effects are largely mediated by activation of the serotonin 5-HT_2A_ receptor in humans[11], while in preclinical research hallucinogenic activity in rodents is operationally modelled by the HTR in rodents[12]. Whether these altered states of consciousness are required for therapeutic efficacy, and the extent to which 5-HT_2A_ receptor engagement contributes to the neuroplastic effects associated with these compounds, remain unresolved questions and continue to be the subject of active investigation[13][14]. Accumulating evidence indicates that, irrespective of the chemical class of hallucinogenic compounds, whether serotonergic psychedelics or dissociative such as ketamine, the neurotrophin brain-derived neurotrophic factor (BDNF) plays a critical role in mediating neuronal plasticity through activation of TrkB receptor[15]. In line with this view, Moliner *et al.* have recently reported that psychedelics promote neuronal plasticity by directly binding to the BDNF receptor TrkB, thereby proposing a central role for TrkB signalling in the plasticity-inducing effects of these compounds[16]. Within the class of serotonergic psychedelics, and beyond their well-documented polypharmacological profiles across multiple G protein-coupled receptors (GPCRs)[17], functional selectivity, or biased agonism, at the 5-HT_2A_ receptor has been described[18]. In this context, several psychedelic analogues display high potency in activating the 5-HT_2A_ receptor while lacking hallucinogenic activity, as exemplified by 2Br-LSD and lisuride in comparison with LSD[12][19] and by ariadne relative to DOI[20]. The molecular mechanisms underlying this biased agonism remain an area of current research efforts and are hypothesized to involve differential engagement of intracellular signaling transducers, including β-arrestins and Gaq proteins, although conclusive evidence supporting this model is currently lacking[21–24]. In the present study, we characterize a previously described murine neural stem cell (NSC)-derived cellular model as an *in vitro* system for the investigation of neuronal plasticity at both morphological and functional levels[25]. To this end, monosynaptic rabies virus-based tracing was implemented to enable quantitative assessment of established synaptic connectivity. In parallel, this cellular platform permits genetic manipulation of NSC-derived lines, allowing for the inducible and selective silencing of genes of interest, including those encoding the 5-HT_2A_ and TrkB receptors.

Within this experimental framework, we examined a panel of classical psychedelics, including tryptamines, phenethylamines, and ergolines, together with their non-hallucinogenic analogues, as well as ketamine and the TrkB-activating ligands brain-derived neurotrophic factor (BDNF) and 7,8-dihydroxyflavone (7,8-DHF). The resulting data indicate that all compounds tested exhibit neuroplastic activity, irrespective of their previously described psychoactive properties. Notably, the facilitation of morphological plasticity, including dendritogenesis, did not display a consistent correspondence with measures of functional plasticity, such as synaptogenesis, nor with the induction of immediate early gene expression. Moreover, selective silencing of 5-HT_2A_ and TrkB receptor expression revealed that the presence of both receptors is required for the induction of neuronal plasticity by serotonergic psychedelics, independent of their hallucinogenic potential. In this sense, the marked attenuation of BDNF-induced neuroplastic effects following 5-HT_2A_ receptor silencing suggests the existence of a functional interaction between 5-HT_2A_ receptor and TrkB-mediated signalling pathways, a conclusion further supported by *in vivo* analyses of dendritic spine density following pharmacological treatment. Recent phosphoproteomic work from our group identified enhanced glycolysis-associated signalling and increased lactate production as features preferentially associated with hallucinogenic serotonergic psychedelics relative to closely related non-hallucinogenic analogues [26]. On this basis, extracellular lactate accumulation was included here as a complementary biochemical readout with which to assess the contribution of 5-HT_2A_ receptor expression and downstream G protein coupling to this metabolic response.

## METHODS

### Drugs

Brain Derived Neurotrophic Factor (BDNF) was obtained from PreproTech (BDNF, 450-02). 7,8-dihydroxyflavone (7,8-DHF, D5446), lysergic acid diethylamide (LSD, L7007), (±)-2,5-Dimethoxy-4-iodoamphetamine hydrochloride (DOI, D101), (±)-ketamine hydrochloride (ketamine, K2753), YM254890 (SML3637) and pertussis toxin (PTX, P7208) were purchased from Sigma-Merck. Lisuride maleate was obtained from Tocris (4052) and 2-bromo-D-lysergic acid diethylamide (2BrLSD, B478840) was purchased from Toronto Research Chemicals (TRC). N,N-Dimethyltryptamine (N,N-DMT, 13959), psilocin (1184) and Ariadne (α-ethyl 2C-D hydrochloride, 26596) were ordered from Cayman Chemical. All drugs were dissolved in dimethyl sulfoxide (DMSO) except for BDNF, ketamine and DOI which were dissolved in water.

### Cell culture and generation of cell lines

Adherent neural stem cell (NSC) lines from the mouse forebrain were obtained following the methodology previously described[27]. Briefly, the fetus dorsal forebrain at E12.5 was dissected and the obtained tissue was disaggregated in order to get a cell suspension. These cells were then transferred to cell-culture plastic plates that had been previously treated with poly-D-lysine 10 μg/ml (Sigma-Merck) to enhance cellular adhesion and promote monolayer cultures. NSCs were maintained and expanded in NS expansion medium composed as follows: DMEM/Ham’s F-12 media with l-glutamine (Gibco), Glucose solution 29mM (Sigma-Merck), MEM nonessential amino acids 1x (Gibco), Penicillin/Streptomycin 1x (Gibco), HEPES buffer solution 4.5 mM (Cytiva), BSA solution 0.012% (Gibco), 2-mercaptoethanol 0.05 mM (Gibco), N2 supplement 1x (Gibco), B27 supplement 1x (Gibco), murine epidermal growth factor (EGF) 10 ng/ml (PeproTech) and human fibroblast growth factor (FGF-2) 10 ng/ml (PeproTech). To induce differentiation of NSCs into various neural cell lineages, the expansion medium was replaced with the following differentiation medium: Neurobasal plus DMEM/Ham’s F-12 media with l-glutamine 1:1 (Gibco), N2 supplement 1x (Gibco), B27 supplement 1x (Gibco), Glutamax 1x (Gibco) and 2-mercaptoethanol 0.05 mM (Gibco). Cells were grown and maintained at 37 °C-5% CO_2_ under a humidified atmosphere. The generation of NSC lines engineered to silence the expression of the *Ntrk2* and *Htr2a* genes was carried out using shRNA technology with lentiviral vectors. Briefly, lentiviral vectors containing the sequences 5’-CCTTAAGGATAACGAACATTTCTCGAGAAATGTTCGTTATCCTTAAGG-3’ and 5’-CCAGGGTGTGTGAACAAGTTTCTCGAGAAACTTGTTCACACACCCTGG-3’, corresponding to *Ntrk2* and *Htr2a* genes, respectively, were ordered from VectorBuilder. These vectors also included a *lac* operon to regulate the transcription of these sequences in an inducible manner upon activation with IPTG (lsopropyl β-D-1-thiogalactopyranoside), along with a puromycin resistance gene. Recombinant lentivirus production was performed following standard Addgene protocols using plasmids containing the shRNA sequences in combination with psPAX2 and pCAG-VSVG as packaging and envelope helper plasmids, respectively. Once recombinant lentiviruses were obtained, NSCs were transfected, and cells surviving puromycin (Invivogen) selection (0.3 µM/mL) were collected and tested for TrkB or 5-HT_2A_ receptor expression after being cultured in IPTG (Thermofisher) containing medium (10 mM). Similarly, we generated an NSC line that permanently expresses the TVA receptor, along with EGFP and rabies virus glycoprotein. In this case, we used the lentiviral pBOB-synP-HTB (30195) plasmid from Addgene, after making a slight modification to remove the sequence corresponding to the H2B human histone. This modification was performed after observing a distinct morphological phenotype in NSCs expressing the unmodified plasmid, which was accompanied by an inability to differentiate into neurons and glia (Supplementary Fig. 1). After transfection with the final recombinant lentivirus, NSCs containing TVA and rabies glycoprotein in their genome were selected by fluorescence-activated cell sorting (FACS) based on EGFP signal with a FACSAriaIII device (Becton Dickinson).

### Immunocytochemistry

Cells were seeded on glass-coverslips pretreated with poly-D-lysine 50 μg/ml (Sigma-Merck) plus laminin 3.3 μg/ml (Sigma-Merck) and fixed with 4% paraformaldehyde before immunolabeling. Primary antibodies used were anti-MAP2 1:200 (Sigma-Merck, M3696), anti-GFAP 1:100 (Cell Signaling Technology, #3670), anti-TrkB 1:100 (R&D Systems AF1494) and anti-5-HT_2A_ 1:100 (Immunostar #242800). Secondary antibodies were anti-mouse IgGAlexaFluor 594 1:1000 (Abcam, ab150105), anti-rabbit IgGAlexaFluor 594 1:1000 (Abcam, ab150076), and anti-goat IgGAlexaFluor 488 1:1000 (Abcam, ab150129). Fluorescent cell nuclei staining was performed with Hoechst 33342 (Invitrogen) according to the manufacturer instructions. Fluorescent images of cells were acquired using a Leica TCS SP5 confocal microscope. Image processing was conducted with Image J software.

### Immunoblotting

Samples were denatured by incubation at 65 °C for 15 minutes and subsequently analyzed via SDS-polyacrylamide gel electrophoresis (SDS-PAGE) using 4-12% bis-Tris gels (NuPAGE, Invitrogen) with MOPS buffer as the running medium. Following separation, proteins were transferred onto nitrocellulose membranes. To minimize nonspecific binding, membranes were blocked for 2 hours at room temperature on a rotating platform in Tris-buffered saline containing 5% non-fat milk and 0.1% Tween-20. The membranes were then incubated overnight with either rabbit polyclonal anti-5-HT_2A_ antibody (1:500, Immunostar #242800) or goat polyclonal anti-TrkB antibody (1:1,000; R&D Systems AF1494). Antibody-protein complexes were detected using horseradish peroxidase-conjugated anti-rabbit IgG (1:5,000; Abcam, ab6721) and anti-goat IgG (1:10,000; Promega #V805A), and immunoreactive bands were visualized by enhanced chemiluminescence (ECL; Cytiva). To verify equal protein loading, membranes were subsequently stripped and reprobed with anti-actin antibody (1:5,000; Hypermol, clone 262).

### Dendritogenesis analysis

Neural stem cells (NSCs) were plated into 12-well plates (120,000 cells/well) pre-coated with poly-D-lysine (10 µg/ml) and laminin (3.3 µg/ml). Following five days of differentiation, dendritic arborization complexity was analyzed using phase-contrast microscopy images at 40x magnification, examining a population of 20 neurons per condition across five distinct microscopic fields. Dendritic complexity was quantified using the Sholl analysis tool in ImageJ software, together with the SNT (Single Neurite Tracker) plug-in to visualize dendritic arborization. This method involves drawing concentric circles centered on the neuronal soma and counting the number of intersections between these circles and the extending dendritic processes. The data were then visualized in a graph representing the number of intersections as a function of distance from the soma. To compare results across experiments, the area under the curve (AUC) was calculated from the center of the soma up to 80 µm. In treatment experiments, pharmacological agents were introduced on the third day of differentiation, followed by the replacement of the culture medium with fresh medium 24 hours post-administration.

### Synaptogenesis analysis

The generation of genetically engineered rabies virus was performed following the protocol previously described[28]. Plasmids pSADdeltaG-F3 (32634), pcDNA-SADB19N (32630), pcDNA-SADB19P (32631), pcDNA-SADB19L (32632), pcDNA-SADB19G (32633) were acquired from Addgene. The original pSADdeltaG-F3 plasmid was modified by subcloning the encoding sequence of mCherry protein between NheI and SacII restriction sites. HEK 293T-TVA 800, B7GG and BHK-EnvA cell lines were generously provided by Edward Callaway from The Salk Institute for Biological Studies (California, USA). The generated NSC line permanently expressing the TVA receptor along with rabies glycoprotein and EGFP was used to obtain neurons in culture that are susceptible to infection by the recombinant and pseudotyped rabies virus. These neurons serve as postsynaptic neurons in this experimental model, as the rabies virus propagates through synapses in a retrograde manner. The optimal ratio of TVA-positive (postsynaptic) cells to TVA-negative (presynaptic) cells required to achieve a measurable difference in synaptogenesis was determined experimentally (Supplementary Fig. 2). The findings established that 20% of the total plated cells should be TVA-positive. This cellular proportion was subsequently employed in all further experiments. To visualize monosynaptic tracing with rabies virus in cultured neurons, a mixed population of NSCs comprising TVA positive cells and those that would give rise to presynaptic neurons was prepared using the total cell suspension and subsequently distributed across 12 well tissue culture plates pretreated with poly-D-lysine (10 µg/mL) and laminin (3.3 µg/mL) at a density of 120,000 cells per well. Following the initiation of differentiation (day 0), cells were exposed to various drug treatments on day 3, infected with recombinant rabies virus on day 5, and subjected to live cell imaging on day 7. Imaging was performed using a Leica DMI8 epifluorescence microscope with a 10x objective after staining cell nuclei with Hoechst 33342. Cells were illuminated to visualize fluorescent signals corresponding to the emission wavelengths of mCherry and Hoechst 33342, which appear red and blue, respectively. Consequently, the blue channel represented the total cell population within the microscopy field, while the red channel identified cells infected by the rabies virus. Ten microscopic field were acquired per well/condition in order to proceed to image analysis. Image analysis for quantifying different cell populations was conducted using FIJI software, utilizing a customized macro. This macro incorporated Gaussian blur sigma filtering before image segmentation, which was based on the autothresholding of cell nuclei stained with Hoechst 33342. Several morphological parameters were adjustable to optimize the segmentation process, including minimum and maximum cell size, minimum circularity, and mCherry fluorescence intensity thresholds (https://github.com/LopezGimenezLab/fiji-macro-synaptogenesis). To assess the extent of synapse formation among neurons within the analyzed microscopy field, the following ratio was used: (100 / blue objects) × red objects = % of mCherry+ cells. About 10.000-15,000 cells were counted per well.

### Immediate early gene expression and RT-qPCR

NSCs were seeded in 12-well tissue culture plates that had been pre-treated with poly-D-lysine (10 µg/mL) and laminin (3.3 µg/mL) at a density of 120,000 cells per well. On the fifth day of differentiation, cells were treated with the various drugs for 45 minutes. Treatment was terminated by placing the plates on ice, and total RNA was extracted from the cells using TRIzol™ Reagent (Invitrogen) in accordance with the protocol provided by the manufacturer. Approximately 500 ng of RNA was reverse transcribed using PrimeScript™ RT Master Mix (Perfect Real Time) (Takara), following the protocol from the manufacturer. Quantification of endogenous transcripts through RT-qPCR was performed using iQ SYBR Green Supermix (Bio-Rad Laboratories) and the CFX Connect Real-Time PCR Detection System (Bio-Rad Laboratories). The following oligonucleotides were used as primers: 5-HT_2A_-Fw, 5’-GCAGTCCATAGCAATGAGC-3’ and 5-HT_2A_-Rv, 5’-GCAGTGGCTTTCTGTTCTCC-3’; TrkB-Fw, 5’-AAGGACTTTCATCGGGAAGC-3’ and TrkB-Rv, 5’-TCGCCCTCCACACAGACAC-3’; c-fos-Fw, 5’-TTCCTGGCAATAGCGTGTTC-3’ and c-fos-Rv, 5’-TTCAGACCACCTCGACAATG-3’; egr-2-Fw, 5’-TGTTAACAGGGTCTGCATGTG-3’ and egr-2-Rv, 5’-AGCGGCAGTGACATTGAAG-3’; bdnf-Fw, 5’-GGCTGACACTTTTGAGCACGTC-3’ and bdnf-Rv 5’-CTCCAAAGGCACTTGACTGCTG-3’. GAPDH served as the internal control gene and was amplified with the oligonucleotides GAPDH-Fw (5’-TGCGACTTCAACAGCAACTC-3’) and GAPDH-Rv (5’-CTTGCTCAGTGTCCTTGCTG-3’). Each transcript in every sample was analyzed in quadruplicate, and the median threshold cycle (CT) value was used to calculate fold change values.

### Cell treatments

The experimental design for the cellular *in vitro* treatments conducted to investigate dendritogenesis, synaptogenesis, lactate accumulation and immediate-early gene expression is summarized in Supplementary Fig. 3. All pharmacological agents were applied to the cell culture medium at a final concentration of 10^-5^ M, except for BDNF, which was administered at 50 ng/mL. Pretreatments to uncouple G_q/11_ and G_i/o_ proteins were performed with YM254890 at 10^-7^M for 2h and PTX at 100ng/mL for 24 h respectively. Drug treatments were removed after 24 h by replacing the culture medium with fresh medium, except in the immediate-early gene expression assays, where treatments were applied for 45 min and lactate accumulation where treatments were applied for 8 h.

### Animals

Experiments were performed on adult (8-14 week-old) C57BL/6J (Jackson Laboratory) male mice. For assays with *5-HT_2A_R* (*Htr2a*) *KO* mice[12], heterozygous mice on a C57BL/6J background were bred to obtain WT and *KO* mice, which were genotyped using tail snips obtained at weaning. All mice were group-housed (up to 5 mice/cage) in non-aseptic individually ventilated cages with a 12-hour light/dark cycle at 23 °C with food and water ad libitum. All experiments were performed during the light cycle. All experiments were conducted in accordance with NIH guidelines and were approved by the Virginia Commonwealth University Institutional Animal Care and Use Committee. All efforts were made to minimize animal suffering and the number of animals used in these experiments.

### In-vivo treatment

The TrkB receptor agonist 7,8-dihydroxyflavone (DHF) was obtained from Abcam (ab120996) and dissolved at 1 mg/ml in 0.9% sterile saline containing 17.5% DMSO. The vehicle-control solution consisted of 0.9% sterile saline with 17.5% DMSO. All animals received intraperitoneal (i.p.) injections at a volume of 5 µl/g, corresponding to a final dose of 5 mg/kg DHF in the experimental group. This dose was selected based on previous studies evaluating the behavioral effects of chronic DHF treatment in mice[29].

Mice received a single daily injection (i.p.) of DHF (5 mg/kg) or vehicle for 14 consecutive days. Samples were collected either 24 h or 48 h after the final DHF or vehicle administration. Because the two vehicle-treated control groups (samples collected 24 h or 48 h after the last vehicle injection) showed no statistically significant differences in frontal cortex dendritic spine density (two-tailed unpaired Student’s t-test: t_61_= 1.31, p > 0.05), these groups were combined into a single vehicle-treated control group for all statistical comparisons with DHF-treated mice.

### Stereotaxic surgery

Mice were anesthetized with isoflurane (2% in 0.5 l/min oxygen, 1% vaporizer setting) in an induction chamber. After induction, the head was shaved around the anticipated surgical site using an electric razor. Mice were then positioned in the stereotaxic frame (Kopf Instruments), secured into the bite bar, and allowed to stabilize under isoflurane delivered through a nose cone (proper breathing rate, negative toe-pinch response) before placement of the ear bars. Anesthesia was adjusted between 1.5-2.5% depending on the mouse’s weight, breathing rate, and toe-pinch response. Toe-pinch checks were recorded every 15 min or before any potentially painful step. A lubricated rectal thermometer connected to a heating pad (Kent Scientific, RightTemp Jr) was inserted to maintain body temperature at 37 °C throughout the procedure. Ophthalmic ointment (Opthavet) was applied to the eyes. The surgical site was disinfected using three alternating swabs of povidone-iodine (PDI) and sterile isopropyl alcohol pads (Dynarex). A sterile scalpel was used to create a midline incision, skin flaps were retracted with bulldog clamps (Fine Science Tools), and the skull was cleaned of membranous tissue with a sterile cotton swab. Bregma was identified, and Hamilton syringes (Kopf, Model 942) were zeroed at this location before moving to the injection sites. Injection coordinates were marked with an autoclaved pencil, followed by bilateral drilling using an Ideal Micro-Drill. Hamilton syringes were lowered bilaterally into the brain at a 10° lateral angle to the target coordinates. Mice of both genotypes (WT and KO) received bilateral injections of 0.3 µl AAV8-CaMKIIa-eYFP-WPRE (UNC Chapel Hill Vector Core) delivered at 0.1 µl/min. The stereotaxic coordinates (relative to bregma and dura) were: +1.6 mm rostrocaudal, - 1 mm dorsoventral, and +2.6 mm mediolateral, with a 10° lateral angle. Following injection, syringes remained in place for 5 min to allow viral diffusion before being slowly withdrawn. Drill holes were sealed with sterile bone wax (Medline), and the incision was closed using Vetbond tissue adhesive (3M). The 14-day DHF/vehicle treatment (see above) began exactly 1 week after surgery so that the treatment paradigm ended 3 weeks post-surgery, when transgene expression is maximal[30]. After DHF/vehicle treatment, mice were deeply anesthetized and transcardially perfused with 20 ml of 1× PBS followed by 20 ml of 4% paraformaldehyde (PFA). Brains were dissected and post-fixed in 4% PFA at 4 °C for 24 h, then transferred to 1× PBS. Coronal sections (40 µm) containing the frontal cortex (including the injection site) were collected using a Leica VT1000S vibratome. Sections were mounted on glass slides (Fisherbrand Premium Superfrost) and coverslipped (Epredia 24 × 50 mm) with ProLong Diamond Antifade Mountant (Invitrogen).

### Dendritic spine analysis

Dendritic spine analysis assays were carried out as previously reported[30], with minor modifications. Briefly, dendrites 50-150 µm away from the soma were randomly selected from eYFP-labeled pyramidal neurons, which were identified by their characteristic triangular soma. Dendritic segments selected for analysis met the following criteria: (i) the segment was completely filled (endings were excluded); (ii) the segment was at least 20 µm from the soma; and (iii) the segment did not overlap with other dendritic branches. Images of pyramidal neurons in layers II/III of the frontal cortex were acquired using confocal microscopy (63-100×, Z-stacks), and dendritic spines were manually counted by a blinded scorer in 20-µm segments.

### Lactate accumulation assay

NSC shRNA 5-HT_2A_ were seeded in 24-well plates at a density of 80.000 cells per well. Following five days of differentiation, cells were treated with the different drugs for 8 h at 37°C and 5% CO_2_ before supernatant collection. These supernatants were centrifuged at 3000g to pellet possible cells detached from the plate during supernatant collection. Supernatants were transferred to sterile microcentrifuge tubes and frozen at -80°C for a period of at least 24 hours. L-lactate concentration was assessed using a modified previous described protocol [31] . Once all independent replicates had been collected, supernatants were thawed, thoroughly mixed, and 20 µl per sample were added to a 96-well plate where they were mixed with 180 µl of a master mix composed by: 14.8 ml of 0.1M Glycine buffer at pH 9.2, 2 ml of 20mM NAD^+^, 2 ml of MTT at 5 mg/ml, 0.6 ml of LDH at 100 U/ml, 0.4 ml of diaphorase at 20 U/ml and 0.2 ml of PMS at 1 mg/ml. The plate was incubated for 30 minutes at 37 °C and absorbance was measured at a wavelength of 570 nm.

### Statistical analysis

All statistical analyses were performed using GraphPad Prism version 10.4. For Sholl analysis, immediate early gene expression, synaptogenesis and lactate accumulation assays data distributions were first assessed for normality using the Shapiro-Wilk test. Expression levels of the immediate early genes *c-fos* and *egr-2* as well as the AUC obtained from Sholl analysis did not conform to a normal distribution; accordingly, these data were log_2_-transformed to permit subsequent parametric statistical analyses. Differences in receptor expression were evaluated using a two-way analysis of variance (ANOVA), with receptor type and silencing condition included as independent factors. The dependent variables were the expression levels of *Htr2a* and *Ntrk2*, normalised to the corresponding non-silenced control condition. Post hoc comparisons were conducted using Fisher’s least significant difference (LSD) test without correction for multiple comparisons. To assess differences in immediate early gene expression, a two-way ANOVA was performed with treatment and silencing condition as independent factors. In data obtained from RT-qPCR experiments, the dependent variable for *bdnf* transcription was normalised to the vehicle-treated condition, whereas for *c-fos* and *egr-2* the dependent variable was the log_2_-transformed normalised expression. Post hoc analyses were conducted using Tukey’s multiple comparisons test. Differences in dendritogenesis and synaptogenesis were analysed using two-way ANOVA, with psychedelic treatment and gene silencing as independent factors. The dependent variables were the area under the curve derived from Sholl intersection analyses and the percentage of total cells expressing mCherry, respectively. Tukey’s multiple comparisons test was applied for post hoc analyses. To evaluate the interaction between 5-HT_2A_ and TrkB signalling *in vivo*, a two-way ANOVA was performed with 7,8-dihydroxyflavone (7,8-DHF) treatment and mouse genotype as independent factors. The dependent variable was dendritic spine density quantified in 20 µm dendritic segments. Post hoc comparisons were performed using Tukey’s multiple comparisons test. To evaluate differences in lactate accumulation experiments, a two-way ANOVA was performed with drug treatment and gene silencing as the independent factors. The dependent variable was the lactate concentration expressed as fold change over vehicle. A follow up Tukey’s multiple comparison test was performed. All data are presented as mean ± standard error of the mean (SEM). Statistical significance was defined as *p* < 0.05. All the experiments were conducted at least three independent times.

## RESULTS

### Development of genetically engineered adherent murine neural stem cell lines

Building upon the neural stem cell (NSC) experimental model originally described by Pollard et al.[27], we recently adapted this approach to facilitate in vitro studies on neuronal plasticity[25]. These neural progenitor cells exhibit a high proliferative capacity in their undifferentiated state and possess the potential to differentiate into various neural lineages, primarily neurons and glial cells. Furthermore, upon differentiation, we observed that the majority of the neuronal population displays a predominantly glutamatergic neurochemical phenotype[25]. In the present study, we investigated the expression of the neurotrophin receptor TrkB and the serotonin 5-HT_2A_ receptor in these cell lines. After five days of differentiation, immunoblotting and immunocytochemistry confirmed the expression of both receptors in neurons and glial cells (Fig. 1a). Notably, TrkB expression increased significantly during differentiation compared to neural progenitors, as demonstrated by immunoblotting (Fig. 1b). In addition, simultaneous immunolabelling experiments demonstrated the co-expression of both receptor in differentiated neurons and glial cells. (Fig. 1a).

**Fig. 1.**
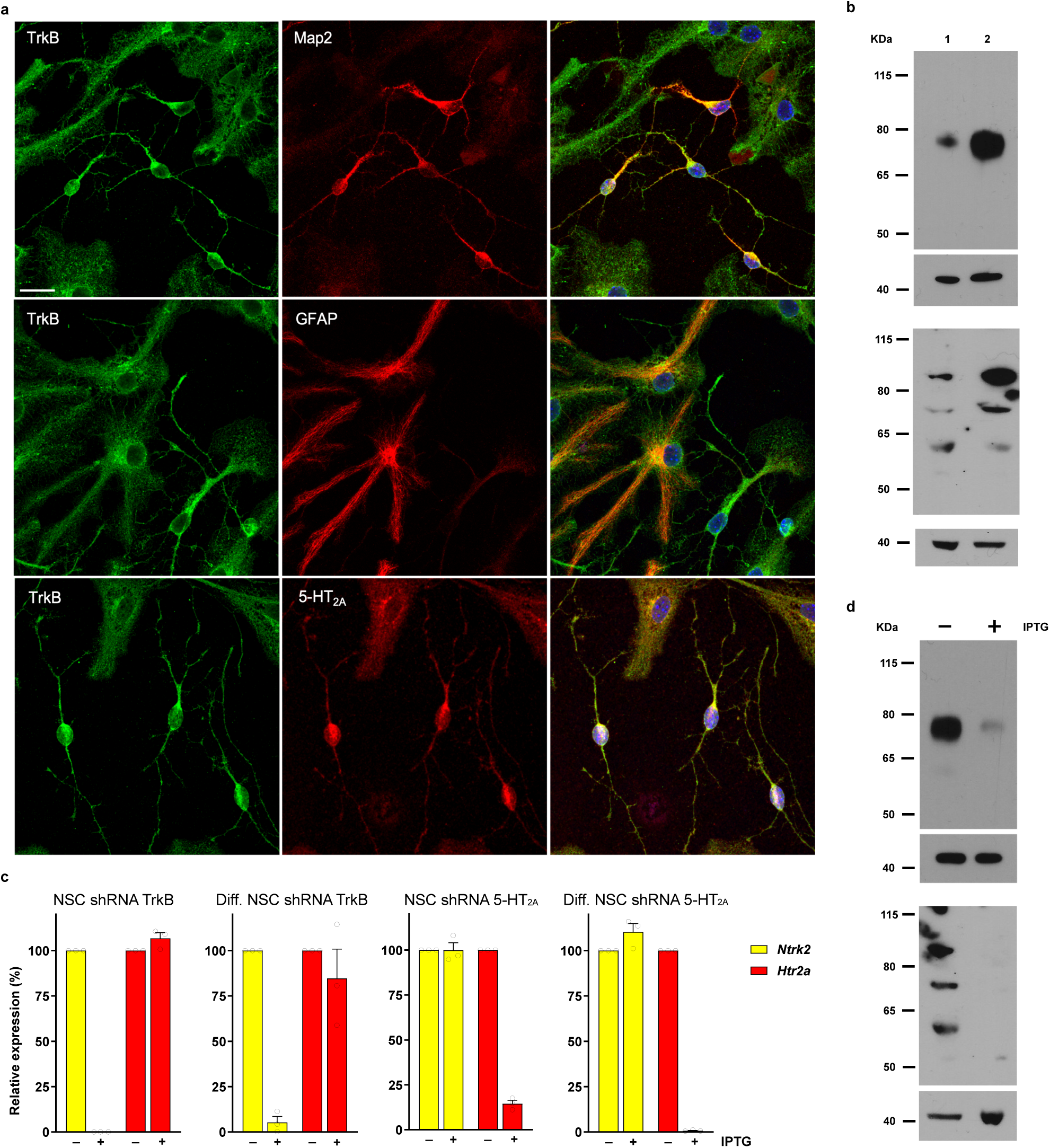
Expression of TrkB and 5-HT_2A_ receptors in neurons and glial cells derived from differentiation of neural stem cells. **a.** Fluorescence microscopy images from immunocytochemistry assays showing the distribution of TrkB receptors (left column) in neural cells differentiated from neural stem cells. The upper row demonstrates co-expression of TrkB receptors with the neuronal marker MAP2, whereas the middle row shows the presence of TrkB receptors in cells expressing the glial marker GFAP. The lower row shows co-expression of TrkB and 5-HT_2A_ receptors in both neurons and glial cells. Scale bar: 20 µm. **b.** Immunoblots displaying protein bands corresponding to TrkB receptors (top) and 5-HT_2A_ receptors (bottom). Lane 1 shows undifferentiated neural stem cells and lane 2 shows neural cells differentiated for 5 days. **c.** Effect of IPTG treatment on the expression of *Ntrk2* and *Htr2a* mRNAs in non-differentiated NSC shRNA 5-HT_2A_, differentiated NSC shRNA 5-HT_2A_, non-differentiated NSC shRNA TrkB and differentiated NSC shRNA TrkB, as determined by RT-qPCR. **d.** Immunoblots showing the effect of IPTG treatment on the expression of TrkB receptors (top) and 5-HT_2A_ receptors (bottom) in cell lysates from NSC shRNA TrkB and NSC shRNA 5-HT_2A_ lines differentiated for 5 days. Immunoblots showing 40 KDa bands correspond to loading controls made with actin immunoreactivity.

To achieve conditional silencing of either TrkB or 5-HT_2A_ receptor expression in these cell lines, we designed shRNA sequences complementary to the mRNA transcripts encoding each receptor. These shRNA constructs were placed under the control of a lac operon promoter and linked to a puromycin resistance gene to facilitate selection. Following lentiviral transfection, NSCs were subjected to antibiotic selection, after which surviving clones were pooled and analyzed for receptor expression. Receptor expression was assessed after 4 days of IPTG treatment in undifferentiated NSCs, and after 5 days of differentiation into neurons and glial cells. Quantitative PCR analyses revealed a robust reduction in *Ntrk2* transcript levels, exceeding 99%, in the NSC shTrkB line (Fig. 1c). Similarly, the NSC sh5-HT_2A_ line exhibited an approximately 85% reduction in *Htr2a* transcript levels (Fig. 1c). In both cases, gene silencing was sustained following differentiation (Fig. 1c), as confirmed by RT-PCR quantification and immunoblotting analyses (Fig. 1d). As a control for shRNA silencing specificity, the expression of the *Ntrk2* and *Htr2a* genes was assessed in cell lines not harbouring their corresponding shRNAs. Quantitative PCR analysis confirmed that each shRNA selectively silenced its respective target gene, with no detectable effect of the transcription of one gene on the transcription of the other (Fig. 1c).

Analogously, for subsequent in vitro synaptogenesis experiments using rabies virus technology, we generated an NSC line stably expressing the TVA receptor, along with EGFP and the rabies virus glycoprotein. Positive clones carrying the recombinant plasmid were selected after lentiviral transfection by cell sorting based on GFP fluorescence. (Supplementary Fig. 2.).

### TrkB and 5-HT_2A_ receptors are essential for neuroplasticity induced by psychedelics

Serotonergic psychedelics, TrkB receptor agonists, and ketamine have all been demonstrated to enhance neuronal morphological plasticity in *in vitro* models, predominantly in primary neuronal cultures derived from mice[4]. In the case of serotonergic psychedelics, which exert their effects at least in part through activation of 5-HT_2A_ receptors, it remains unresolved whether 5-HT_2A_ receptor stimulation is indispensable for dendritogenesis, or whether this process is mediated exclusively through TrkB receptor signaling, as has been suggested in recent studies[16]. To validate our neuronal culture system derived from differentiated neural progenitors, we evaluated dendritic arborization by means of Sholl analysis. Cells were exposed to the compounds for 24 hours, and imaging was performed 48 hours post-treatment. All drugs tested induced a significant increase in dendritic complexity relative to basal conditions (Fig. 2a). Silencing of either TrkB or 5-HT_2A_ receptors did not alter basal dendritic complexity under vehicle conditions (Supplementary Fig. 4 and Supplementary Fig. 7). However, suppression of TrkB expression abolished the dendritogenic response to all compounds examined (Fig. 2a). Similarly, silencing of 5-HT_2A_ receptors eliminated the effects of serotonergic psychedelics (DMT, PSI, DOI) and non-psychedelics (Ariadne, 2-Br-LSD and Lisuride). In contrast, BDNF, 7,8-DHF, and ketamine remained partially effective, although their activity was attenuated compared with cells expressing 5-HT_2A_ receptors (Fig. 2a). The reduced responsiveness to TrkB agonists following 5-HT_2A_ ablation suggested a mechanism involving decreased BDNF synthesis. To address this hypothesis, we measured *Bdnf* transcript levels after 45 minutes of drug exposure. Quantitative PCR analysis revealed that TrkB agonists (BDNF and 7,8-DHF), tryptamines (PSI and DMT) and ergolines (LSD, 2-Br-LSD and Lisuride) but not ketamine and phenethylamines (DOI and Ariadne) increased *Bdnf* expression (Supplementary Fig. 5). This effect was entirely abolished by silencing of 5-HT_2A_ receptors (Supplementary Fig. 5a), whereas basal *Bdnf* transcription remained unaltered (Supplementary Fig. 5b).

**Fig. 2.**
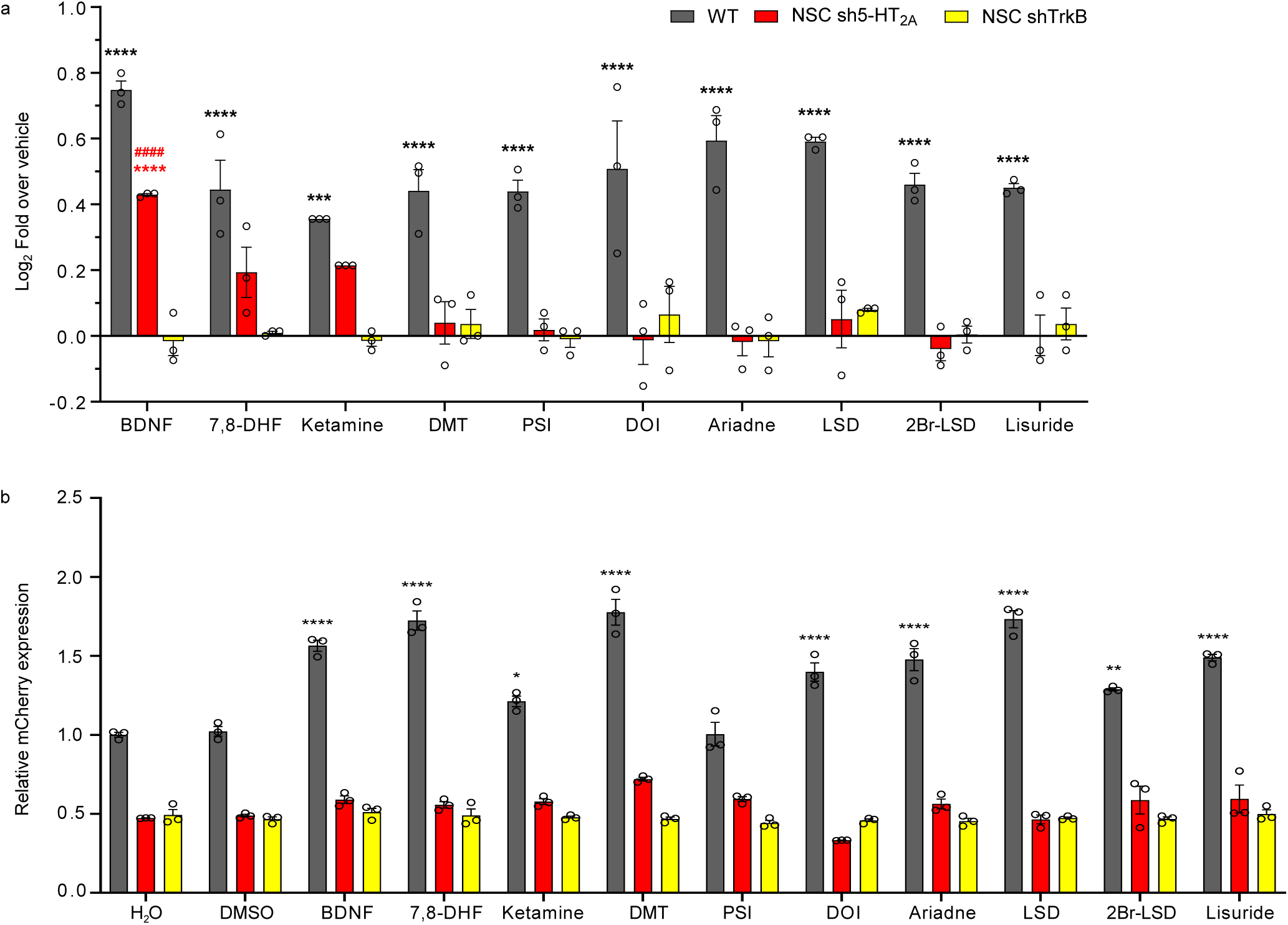
TrkB and 5HT_2A_ receptors drive dendritogenesis induced by psychedelics. **a.** Wild-type NSCs (grey bars), TrkB-silenced NSCs (NSC shRNA TrkB, yellow bars), and 5-HT_2A_-silenced NSCs (NSC shRNA 5-HT_2A_, red bars) were differentiated into neurons and glia and treated with the indicated drugs. Bars represent the fold change of the area under the curve (AUC) obtained from Sholl analysis. Two-way ANOVA revealed significant main effects of phenotype (F(2,60) = 226.600, p < 0.0001) and treatment (F(9,60) = 9.614, p < 0.0001), as well as a significant interaction (F(18,60) = 5.620, p < 0.0001). Post hoc comparisons were performed using Tukey’s multiple comparison test. *p < 0.05, **p < 0.01, ***p < 0.001, ****p < 0.0001. # indicates significant differences between WT and NSC shRNA TrkB. Same p values are applied. **b.** Cell mixtures of NSC TVA with wild- type NSCs (grey bars), TrkB-silenced NSCs (NSC shRNA TrkB, yellow bars), and 5-HT_2A_-silenced NSCs (NSC shRNA 5-HT_2A_, red bars) were differentiated into neurons and glia and treated with the indicated drugs. Bars represent the relative number of mCherry-positive cells versus vehicle. Statistical analysis was performed using two-way ANOVA, which revealed significant effects of phenotype (F(2,72) = 2008, p < 0.0001), drug treatment (F(11,72) = 23.44, p < 0.0001), and their interaction (F(22,72) = 17.81, p < 0.0001). Post hoc comparisons were performed using Tukey’s multiple comparison test. *p < 0.05, **p < 0.01, ***p < 0.001, ****p < 0.0001. # indicates significant differences between the silenced condition and WT among applying vehicles.

To further explore whether the morphological neuroplasticity induced by drug treatments is accompanied by corresponding changes in functional neuronal plasticity, such as synaptic connectivity, we adapted a rabies virus-based methodology originally developed for neuronal tracing in intact brain tissue [32, 33]. For this purpose, we established a neural stem cell (NSC) line that constitutively expressed the TVA receptor together with EGFP and the rabies glycoprotein. Upon differentiation, these cells became susceptible to infection by a rabies virus engineered to express the fluorescent protein mCherry and pseudotyped with EnvA glycoprotein for selective interaction with TVA receptors. The TVA positive neurons served as starter cells, enabling retrograde trans-synaptic spread of the rabies virus to co-cultured neurons that did not express TVA. In this system, neurons differentiated from NSC lines containing either shTrkB or sh5HT_2A_ sequences acted as presynaptic partners of the TVA positive cells (Supplementary Fig. 2a). Following drug treatments, the number of mCherry-positive neurons was quantified, providing a readout of potential drug-induced changes in synaptic connectivity relative to vehicle-treated controls. This analysis revealed a significant increase in the number of neurons containing mCherry-positive rabies virus in all cases, except for psilocin, where no difference was observed relative to the vehicle condition (Fig. 2b). Silencing of TrkB or 5HT_2A_ receptors abolished these drugs induced effects, as equivalent treatments did not produce significant changes in the number of mCherry positive neurons compared with the corresponding vehicle controls. Irrespective of the individual drug effects on synaptogenesis, ablation of TrkB or 5HT_2A_ receptor expression markedly reduced the basal level of neuronal connectivity, as indicated by the comparison of vehicle conditions in receptor silenced cells with those in cells expressing both receptors (Fig. 2b).

To assess whether TrkB-dependent structural plasticity observed *in vitro* was recapitulated *in vivo*, dendritic spine density was quantified in the frontal cortex of mice following reapeated administration of the TrkB agonist 7,8-DHF. Wild-type mice received daily treatment with 7,8-DHF for 14 days, and dendritic spines were analyzed at 24 and 48 hours after the final dose. At 24 hours post-treatment, 7,8-DHF-treated wild-type mice exhibited a significant reduction in dendritic spine density compared with vehicle-treated controls (Supplementary Fig. 6). This reduction was no longer detectable at 48 hours after the final dose, with spine density returning to levels comparable to vehicle-treated animals (Supplementary Fig. 6). The same treatment paradigm was applied to 5-HT_2A_ receptor knockout (5-HT2AR-KO) mice. In contrast to wild-type animals, 7,8-DHF treatment did not produce a significant decrease in dendritic spine density in 5-HT_2A_R-KO mice at the 24-hour time point (Supplementary Fig. 6). At this time point, a non-significant trend toward increased spine density was observed in 7,8-DHF-treated 5-HT_2A_R-KO mice relative to vehicle controls (p = 0.06). No significant differences between treatment groups were detected at 48 hours in 5-HT2AR-KO mice. Together, these results indicate that chronic 7,8-DHF administration produces transient alterations in dendritic spine density in the frontal cortex *in vivo*, and that the presence of the 5-HT_2A_ receptor influences the direction and magnitude of these effects.

### Immediate-early gene expression induced by psychedelics are mediated by TrkB and 5-HT_2A_ receptors

Transcript levels of the immediate early genes *c-Fos* and *Egr-2* were quantified following acute pharmacological stimulation in differentiated neuronal and glial cultures. In cells expressing both TrkB and 5-HT_2A_ receptors, most compounds significantly increased *c-Fos* and *Egr-2* expression relative to vehicle controls. The only exception was ketamine for c-fos induced expression (Fig. 3a). Silencing of either TrkB or 5-HT_2A_ receptors did not significantly alter basal *c-Fos* or *Egr-2* expression under vehicle conditions (H20 or DMSO; Supplementary Fig. 7). In contrast, receptor suppression markedly affected drug-evoked transcriptional responses. TrkB silencing abolished the induction of *c-Fos* and *Egr-2* across all compounds tested, reducing transcript levels to those observed under vehicle conditions. Similarly, 5-HT_2A_ receptor silencing eliminated drug-induced increases in both genes, with the exception of BDNF, which retained the ability to elevate *c-Fos* and *Egr-2* expression, and ketamine, which did not significantly affect *Egr-2* expression (Fig. 3a,b).

**Fig. 3.**
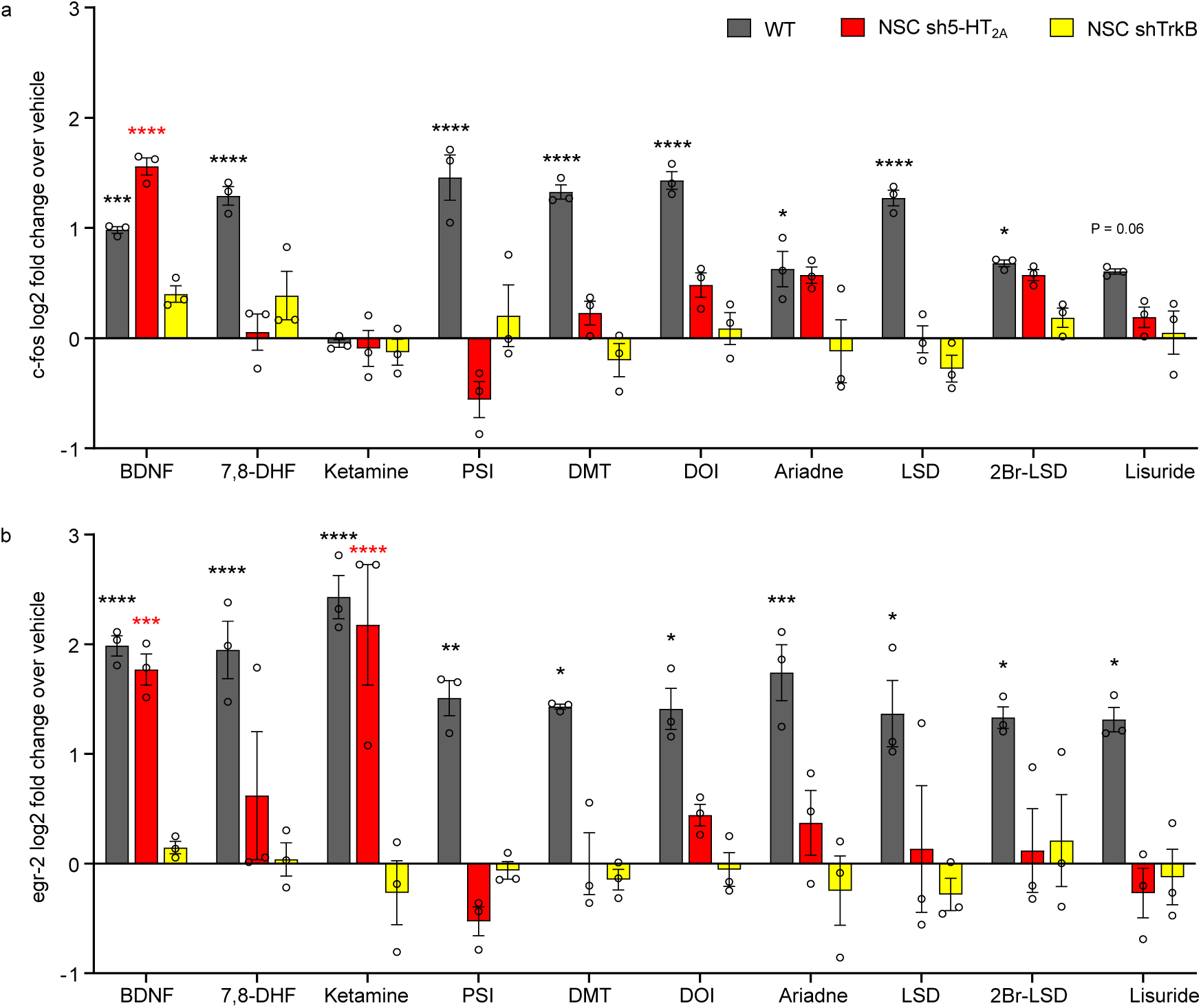
Effect of psychedelic drug treatments on the expression of immediate early genes *c-Fos* and *Egr-2*. Wild-type NSCs (grey bars), TrkB-silenced NSCs (NSC shRNA TrkB, yellow bars), and 5-HT_2A_-silenced NSCs (NSC shRNA 5-HT_2A_, red bars) were differentiated into neurons and glia and treated with the indicated drugs to assess changes in immediate early gene expression. **a.** *c-Fos* expression, represented as log2 fold change relative to vehicle levels. Two-way ANOVA revealed significant effects of drug treatment (F(10,66) = 7.214, p < 0.0001), NSC phenotype (F(2,66) = 50.07, p < 0.0001), and their interaction (F(20,66) = 3.231, p = 0.0002). Post hoc comparisons were performed using Tukey’s multiple comparison test. *p < 0.05, **p < 0.01, ***p < 0.001, ****p < 0.0001. **b.** *Egr-2* expression, represented as log2 fold change relative to vehicle levels. Two-way ANOVA revealed significant effects of drug treatment (F(10,66) = 6.872, p < 0.0001), NSC phenotype (F(2,66) = 85.90, p < 0.0001), and their interaction (F(20,66) = 3.162, p = 0.0002). Post hoc comparisons were performed using Tukey’s multiple comparison test versus vehicle. *p < 0.05, **p < 0.01, ***p < 0.001, ****p < 0.0001.

### Participation of G_q/11_ and G_i/o_ proteins in neuroplasticity and hallucinogenic-like responses elicited by psychedelics

To assess the contribution of G_q/11_ and G_i/o_ proteins as transducing signalling elements in psychedelic actions mediated by GPCRs, pharmacological uncoupling strategies were employed. The selective G_q/11_ inhibitor YM254890 and pertussis toxin (PTX), which functionally uncouples G_i/o_ proteins, were used to evaluate their impact on neuroplastic responses. In parallel, lactate production in the culture medium was measured as an indicator of metabolic activity associated with psychedelic compounds displaying hallucinogenic properties [26].

Analysis of dendritogenesis demonstrated that pretreatment with either YM254890 or PTX attenuated the neuroplasticity-enhancing effects of both psychedelic (DMT, PSI, DOI, LSD) and non-psychedelic (Ariadne, 2Br-LSD, Lisuride) compounds examined (Fig. 4a). LSD represented an exception, as uncoupling of either G_q/11_ or G_i/o_ proteins did not alter this morphological response (Fig. 4a). Functional neuroplasticity was further assessed through synaptogenesis experiments. Pretreatment with G protein uncoupling agents resulted in a marked increase in synapse number relative to basal or vehicle conditions (Fig. 4d). Under G_q/11_ inhibition with YM254890, this elevation in basal synaptic number prevented the detection of any additional facilitatory effects of the psychedelic compounds relative to their basal levels (Fig. 4d). A comparable pattern was observed following G_i/o_ uncoupling with PTX. However, under these conditions treatment with DOI or LSD significantly reduced synapse number relative to their basal controls, reaching levels comparable to those observed under basal conditions in the absence of G protein uncoupling (Fig. 4d).

**Fig. 4.**
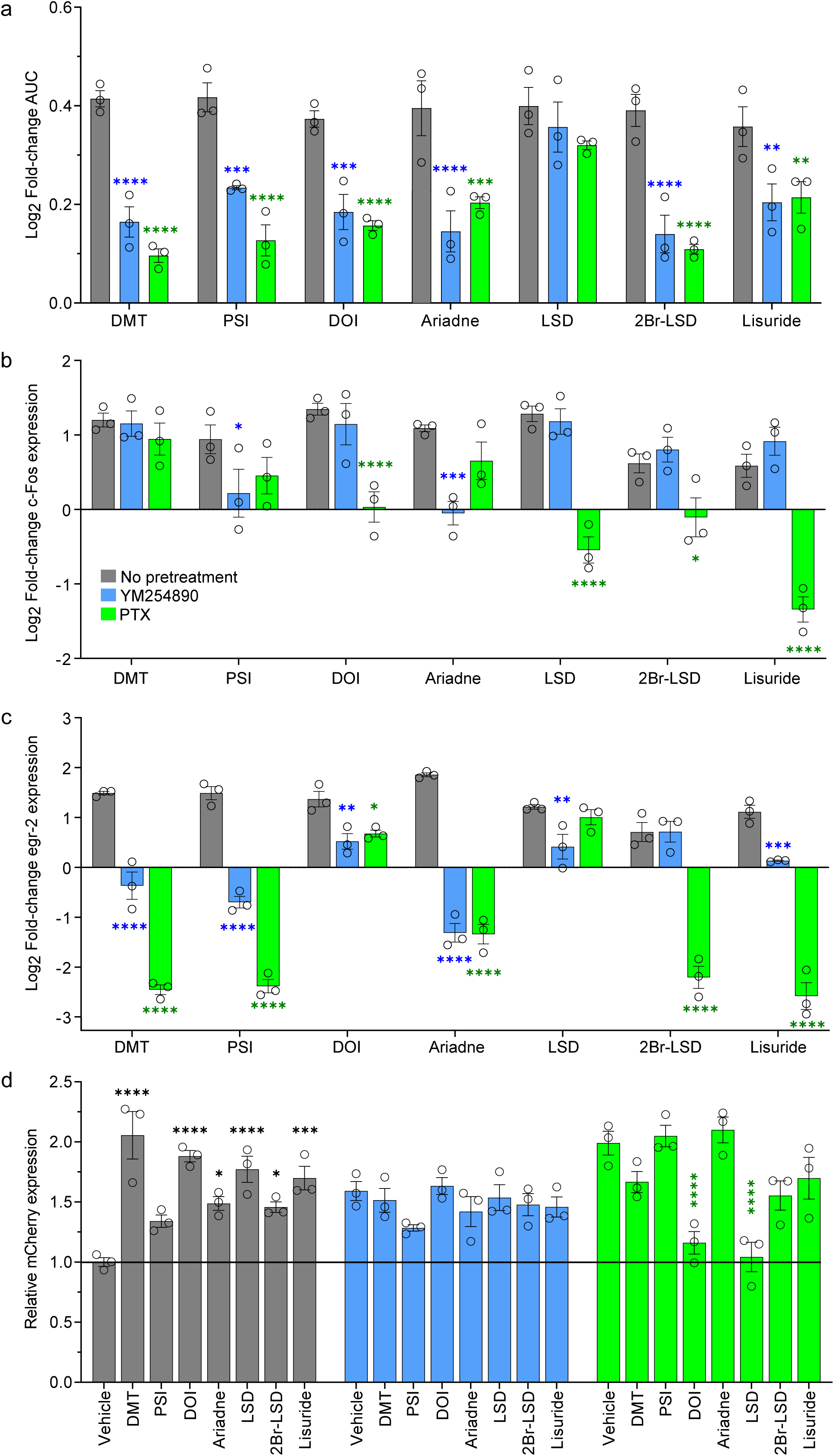
Effect of G protein decoupling in neuroplasticity and immediate early gene expression assays. Wild type NSCs were differentiated into neurons and glia and pretreated with YM254890 (blue bars), PTX (green bars) or vehicle (grey bars); cells were then treated with drugs and dendritogenesis, immediate early gene expression and synaptogenesis was assessed. **a.** Bars represent the log2 fold change of the AUC obtained from Sholl analysis. Two-way ANOVA revealed significant main effects of treatments (F(6,42) = 7.096, p < 0.0001) and pretreatments (F(2,42) = 98.680, p < 0.0001), as well as a significant interaction (F(12,42) = 2.657, p = 0.0096). Post hoc comparisons were performed using Tukey’s multiple comparison test. *p < 0.05, **p < 0.01, ***p < 0.001, ****p < 0.0001. **b.** Bars represent the log2 fold change of the expression of *c-Fos* relativized to its vehicle. Two-way ANOVA revealed significant main effects of treatments (F(6,42) = 8.717, p < 0.0001) and pretreatments (F(2,42) = 51.510, p < 0.0001), as well as a significant interaction (F(12,42) = 8.505, p < 0.0001). Post hoc comparisons were performed using Tukey’s multiple comparison test. *p < 0.05, **p < 0.01, ***p < 0.001, ****p < 0.0001. **c.** Bars represent the log2 fold change of the expression of *Egr-2* relativized to its vehicle. Two-way ANOVA revealed significant main effects of treatments (F(6,42) =42.86, p < 0.0001) and pretreatments (F(2,42) = 455.80, p < 0.0001), as well as a significant interaction (F(12,42) = 34.97, p < 0.0001). Post hoc comparisons were performed using Tukey’s multiple comparison test. *p < 0.05, **p < 0.01, ***p < 0.001, ****p < 0.0001. **d.** Bars represent the relative number of mCherry^+^ cells relativized to the non-pretreated vehicle. One-way ANOVA of non-pretreated cells revealed significant effect of the treatments (F(7,16) = 12.46, p <0.0001). One-way ANOVA of YM254890-pretreated cells revealed no significant effect of the treatments (F(7,16) = 1.45, p = 0.2535). One-way ANOVA of PTX-pretreated cells revealed significant effect of the treatments (F(7,16) = 11.85, p <0.0001).

The effects of G protein uncoupling on immediate early gene expression were next examined. G_q/11_ uncoupling produced only limited effects on *c-Fos* induction (Fig. 4b), with significant changes observed only for psilocin and Ariadne relative to basal levels. In contrast, PTX pretreatment significantly modified the *c-Fos* response to DOI, LSD, 2Br-LSD and lisuride, indicating a greater contribution of G_i/o_ signalling to the regulation of this transcriptional response. A more pronounced effect was observed for *Egr-*2 expression (Fig. 4c). Pretreatment with YM254890 markedly altered psychedelic-induced *Egr-2* responses relative to treatments in the absence of the uncoupling agent, indicating a prominent role for G_q/11_ signalling in this transcriptional regulation. G_i/o_ uncoupling with PTX similarly produced substantial modulation of *Egr-2* expression, with responses significantly altered for most compounds examined. Nevertheless, LSD was the only ligand for which PTX pretreatment did not modify *Egr-2* induction. Collectively, these findings indicate that both G_q/11_ and G_i/o_-mediated signalling pathways contribute to the structural and transcriptional plasticity associated with psychedelic-induced neuronal activation.

We next assessed the effect of psychedelic compounds on lactate production. As previously reported [26], in control cells, the psychedelics DMT, psilocin, DOI and LSD produced a clear increase in extracellular lactate levels, an effect not observed with the non-psychedelic compounds Ariadne, 2Br-LSD or Lisuride (Fig. 5a). In contrast, knockdown of 5-HT_2A_ receptor expression abolished this psychedelic-specific response (Fig. 5a), indicating that lactate production induced by these compounds is dependent on the presence of the 5-HT_2A_ receptor. The contribution of downstream G protein signalling pathways was then examined using pharmacological uncoupling approaches (Fig. 5b).

**Fig. 5.**
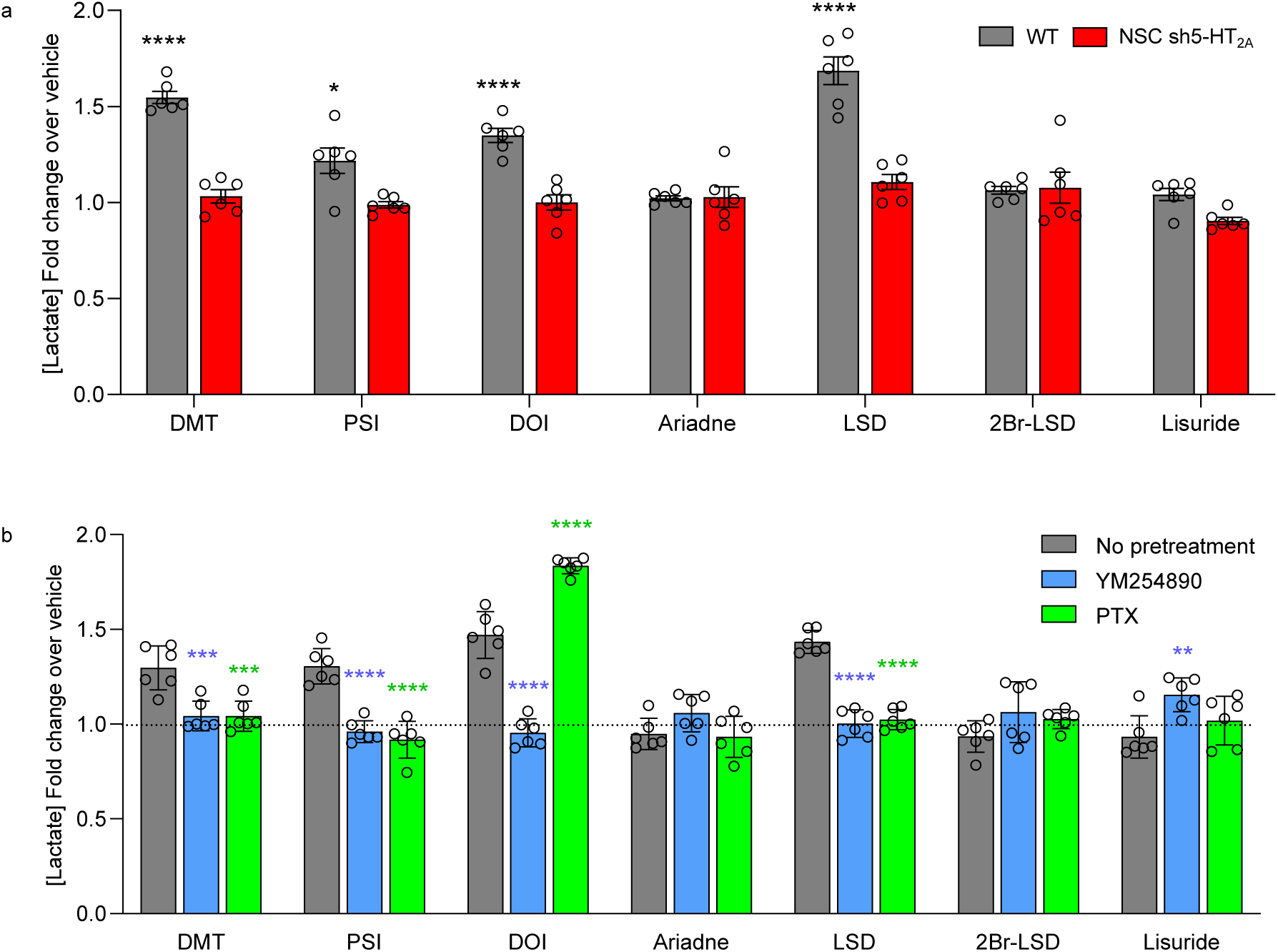
Hallucinogenic serotonergic psychedelics induce lactate accumulation through a 5-HT_2A_ receptor-dependent mechanism. Non-pretreated wild-type NSCs (grey bars), NSCs shRNA 5-HT_2A_ (red bars), YM-254890 pretreated wild-type NSCs (blue bars) and PTX-pretreated wild-type NSCs (green bars) where differentiated into neurons and glia and treated with drugs to assess lactate accumulation in the culture supernatants. **a.** Hallucinogenic psychedelics induce lactate accumulation while non-hallucinogenic analogs lack this property. Silencing 5-HT_2A_ abolishes this response. Two-way ANOVA revealed significant main effects of treatments (F(7,80) =24.90, p < 0.0001) and silencing (F(1,80) = 112.50, p < 0.0001), as well as a significant interaction (F(7,80) = 15.83, p < 0.0001). Post hoc comparisons were performed using Tukey’s multiple comparison test. *p < 0.05, **p < 0.01, ***p < 0.001, ****p < 0.0001. **b.** Decoupling G proteins abolished the response elicited by hallucinogenic psychedelics with the exception of PTX-pretreated DOI treatment which increased. Two-way ANOVA revealed significant main effects of treatments (F(6,105) = 46.19, p < 0.0001) and pretreatment (F(2,105) = 29.22, p < 0.0001), as well as a significant interaction (F(12,105) = 34.74, p < 0.0001). Post hoc comparisons were performed using Tukey’s multiple comparison test. *p < 0.05, **p < 0.01, ***p < 0.001, ****p < 0.0001.

Pretreatment with YM254890 or PTX generally prevented the increase in lactate production elicited by the psychedelic compounds. A notable exception was observed for DOI following PTX pretreatment, where lactate production was not only preserved but significantly potentiated relative to control conditions. Additionally, YM254890 pretreatment led to a weak increase in lactate production in response to Lisuride compared to its respective control group. Together, these observations indicate that 5-HT_2A_ receptor activation is required for the lactate response induced by psychedelics, while also revealing distinct contributions of G_q/11_ and Gi/o protein-dependent signalling pathways in the regulation of this metabolic output.

## Discussion

The renewed interest in serotonergic psychedelics as potential therapeutic approaches for mental health disorders over the past decade has stimulated extensive investigation into the mechanisms underlying their putative therapeutic effects [34]. From a pharmacological perspective, two broad classes of responses can be observed following the administration of psychedelic compounds in humans [3]. The first corresponds to an acute and transient psychoactive response, characterised primarily by altered states of consciousness. The second consists of a more sustained therapeutic effect that emerges following one or a limited number of administrations and persists over the medium to long term. Based largely on preclinical studies, this latter response has been hypothesised to involve the activation of neuronal plasticity-related processes [5,6]. Whether the therapeutic efficacy of psychedelic compounds is associated with both of these responses, or primarily with the neuroplasticity-related component, remains a matter of considerable debate and active investigation [14, 35]. In this context, the availability of psychedelic-derived molecules that lack overt psychoactive properties, at least in preclinical assays of hallucinogenic-like activity such as the head twitch response in mice, provides a valuable opportunity to explore the molecular basis of this apparent functional dissociation. Notable examples include 2-Br-LSD[19] and lisuride[12, 36], which are structurally related to LSD, as well as Ariadne, a derivative of DOI[20]. In the present study, we characterized an *in vitro* cellular model derived from neural stem cells capable of differentiating into neuronal and glial lineages, providing a suitable platform for investigating the molecular pharmacology of psychedelic action. Using this system, we examined several cellular responses associated with psychedelic exposure, including morphological and functional measures of neuronal plasticity, induction of immediate-early gene expression and metabolic regulation reflected by lactate production. These endpoints were assessed in response to a panel of serotonergic psychedelics encompassing representatives of the three principal chemical families of classical psychedelics, namely tryptamines, phenethylamines and ergolines, including both psychedelic compounds and selected non-psychedelic analogues, as well as ketamine. TrkB receptor agonists were included as positive controls for neurotrophic signalling and plasticity-related processes. This cellular model based on neural progenitors is proliferative and self-renewing in its undifferentiated state, providing a continuous source of biological material. As such, it offers a practical alternative to traditional primary cultures and allows a substantial reduction in the use of experimental animals. In addition, the system permits generation of derivative cell lines in which selected genes can be manipulated through targeted silencing or transgenic expression. In previous studies we characterised the neuronal population obtained following differentiation as predominantly glutamatergic [25]. In the present work we further demonstrate endogenous expression of both 5-HT_2A_ and TrkB receptors in neuronal and glial populations within this culture system. Moreover, using RNA interference technology, we generated inducible cell lines in which the genes encoding either the 5-HT_2A_ or TrkB receptors could be selectively silenced. Neuroplasticity assays in this model largely recapitulated findings previously reported in primary murine cultures with regard to the induction of dendritogenesis, as assessed by Sholl analysis [4, 19, 37]. At this morphological endpoint, no significant differences were detected between psychedelic and non-psychedelic serotonergic compounds, as all of the serotonergic ligands examined proved capable of eliciting this response. Comparable effects were also observed with ketamine and with TrkB receptor agonists. Silencing of TrkB expression resulted in complete ablation of the dendritogenic response to all compounds tested, indicating that TrkB is required for the potentiation of neuroplasticity induced by serotonergic psychedelics, ketamine and TrkB agonists, consistent with previous studies employing TrkB antagonists or mutant receptor forms that established a central role for TrkB signalling in psychedelic-induced structural plasticity [4, 16]. By contrast, although silencing of the 5-HT_2A_ receptor also proved essential for the neuroplastic effects of serotonergic psychedelics, it did not completely abolish the response to ketamine, 7,8-DHF or BDNF. Instead, the magnitude of the effect was significantly reduced, with BDNF remaining capable of eliciting a detectable neuroplastic response. These findings suggest that whereas 5-HT_2A_ receptor signalling is necessary for the dendritogenic actions of serotonergic psychedelics, TrkB-mediated mechanisms can under certain conditions, support structural plasticity independently of 5-HT_2A_ receptor activation. At first sight, these results appear to diverge from those reported by Moliner *et al.*, who concluded, on the basis of pharmacological antagonism of the 5-HT_2A_ receptor that the dendritogenic effects of LSD and psilocin are mediated exclusively through TrkB and are independent of 5-HT_2A_ receptor-dependent signalling [16]. However, the requirement for 5-HT_2A_ receptor expression observed here is consistent with other recent reports indicating that serotonergic psychedelics engage 5-HT_2A_ receptor-dependent mechanisms to initiate or facilitate neuroplastic responses [38, 39]. The pharmacological interaction observed between TrkB and 5-HT_2A_ receptors in the *in vitro* neuroplasticity experiments was further examined *in vivo* using 5-HT_2A_ knockout animals. Although the pattern of effects observed *in vivo* differed from those obtained in the 5-HT_2A_ shRNA neural stem cell-derived cultures, where loss of the receptor reduced dendritogenesis and abolished BDNF synthesis and synaptogenesis, the key observation is that the response to TrkB activation remained clearly dependent on the presence or absence of the 5-HT_2A_ receptor. In the *in vivo* experiments, the TrkB agonist 7,8-DHF retained the capacity to promote spinogenesis in the frontal cortex of 5-HT_2A_ knockout animals, suggesting that removal of 5-HT_2A_ receptor signalling alters, rather than abolishes, the downstream consequences of TrkB activation. One proposed mechanism underlying the interaction between 5-HT_2A_ and TrkB receptors is the induction of BDNF synthesis following activation of the 5-HT_2A_ receptor by psychedelic compounds [4, 16, 40]. Consistent with this view, stimulation of 5-HT_2A_ receptors has been reported to increase *Bdnf* transcription in cortical systems, including after administration of the psychedelic 5-HT_2A_ receptor agonist DOI (for an extensive review see [41]). In our model, most compounds tested increased *Bdnf* mRNA transcription; however, this response was not observed for the phenethylamines DOI and Ariadne. One possible explanation is the comparatively higher selectivity of phenethylamines for receptors of the 5-HT_2_ receptor family [3, 17], which may limit the engagement of additional signalling pathways mediated by other G proteins or effector systems. This interpretation is consistent with other observations in the present study, including the distinctive behaviour of DOI in the lactate experiments following G_i/o_ protein uncoupling with PTX, and supports the notion that the functional interaction between 5-HT_2A_ and TrkB signalling may depend on the specific signalling repertoire engaged by individual ligands. Taken together, these findings indicate that the relationship between 5-HT_2A_ and TrkB signalling is unlikely to follow a simple hierarchical organization and instead supports the existence of a functional interaction in which the presence of 5-HT_2A_ receptors modulates the neuroplastic responses mediated by TrkB signalling.

A further innovative aspect of this study is the implementation of modified rabies virus-based synaptic tracing to examine pharmacologically regulated synaptogenesis *in vitro*. This approach, which provides a functional assessment of synaptic connectivity, broadly confirmed previous findings obtained using methods based on the colocalization of pre-and postsynaptic proteins, in that most compounds tested increased the number of synaptic contacts within the neuronal population[4]. At the same time, it revealed several noteworthy features not readily apparent with more conventional approaches. In particular, basal synapse number was markedly reduced following silencing of either 5-HT_2A_ or TrkB receptors, suggesting that both receptors contribute to synapse formation or maintenance under basal conditions, independently of any detectable effect on neuronal viability. Equally notable was the lack of effect of psilocin on synaptogenesis, in contrast to the other serotonergic psychedelics examined. This distinction was not observed in the dendritogenesis experiments, indicating that morphological and functional measures of neuroplasticity do not necessarily correlate fully in this model. This interpretation is further supported by the differential effect of 5-HT_2A_ receptor silencing on the neuroplastic responses induced by TrkB agonists. With regard to psilocin, recent findings showing that this compound, unlike DMT, does not increase sEPSC amplitude also support the possibility that psilocin exerts a distinct functional profile at the synaptic level[42]. When considered together with our phosphoproteomic data (Supplementary Table 1) [26], in which psilocin uniquely regulated a subset of phosphopeptides linked to cytoskeletal proteins, these observations suggest that psilocin may elicit a phenotypic signature in neurons that differs from that of other serotonergic psychedelics.

Immediate-early genes constitute a well-established downstream readout of psychedelic-induced neuronal activation, and both *c-Fos* and *Egr-2* have repeatedly been shown to increase following 5-HT_2A_ receptor stimulation, albeit with ligand-specific transcriptional fingerprints[3]. In particular, González-Maeso *et al.* showed that both LSD and lisuride increased *c-Fos*, whereas induction of *Egr-1* and *Egr-2* was more closely associated with hallucinogenic signalling[12], indicating that these genes may better discriminate between psychedelic and non-psychedelic 5-HT_2A_ receptor agonists. In our model, most serotonergic compounds increased *c-Fos* and *Egr-2* under control conditions, but silencing of either 5-HT_2A_ or TrkB largely abolished these responses. This indicates that immediate-early gene induction in this system depends not only on 5-HT_2A_ receptor activation but also on intact TrkB signalling, and further supports a functional interaction between these receptor systems. By contrast, the responses elicited by ketamine and TrkB agonists are likely to reflect convergence on neuroplasticity-associated transcriptional programmes downstream of BDNF-TrkB signalling, rather than recruitment of the 5-HT_2A_ receptor-dependent hallucinogenic transcriptional signature described for serotonergic psychedelics[43].

In a parallel study using this same neural stem cell-derived model, together with an extensive phosphoproteomic analysis following treatment with this panel of 5-HT_2A_ receptor agonists, we identified a differential signalling signature that distinguished psychedelic from non-psychedelic ligands on the basis of either their psychoactive effects in humans or their activity in the mouse head-twitch response assay [26]. The transductional basis of this functional selectivity remains a matter of considerable debate [22]. One view holds that the hallucinogenic component is primarily dependent on G_q/11_ protein signalling, with differences between psychedelic and non-psychedelic agonists reflecting the efficiency with which the receptor couples to this class of G proteins, whereas induction of neuroplasticity-related processes has been proposed to depend more strongly on β-arrestin-mediated pathways [19, 21, 23]. More recently, however, other studies have suggested that G_i/o_ protein-dependent signalling may make a major contribution to the psychoactive response [44, 45]. To address this question in our system, we performed experiments using YM254890 and PTX to uncouple G_q/11_ and G_i/o_ proteins, respectively. A particular strength of this approach is that these experiments were carried out in differentiated neuronal and glial cultures expressing an endogenous complement of receptors, G proteins and downstream effectors, rather than in heterologous overexpression systems using engineered biosensors or reporter-tagged transducers, as in many previous studies of receptor-G protein coupling. In the morphological neuroplasticity assays, uncoupling of either G_q/11_ or G_i/o_ protein signalling significantly reduced, in most cases, the dendritogenic response induced by serotonergic psychedelics. LSD represented a notable exception, as its effect was not significantly affected by pretreatment with either uncoupling agent. One possible explanation for this divergence lies in the broader pharmacological profile of LSD relative to the other tryptamines and phenethylamines included in the study, which may allow recruitment of additional signalling mechanisms not equivalently engaged by the other ligands[17]. Interpretation of the functional plasticity experiments, assessed through synaptogenesis, is less straightforward, as pretreatment with the G-protein uncoupling agents produced an unexpected elevation in basal synapse number. This increase in baseline connectivity complicates the evaluation of the pharmacologically induced responses and is itself of considerable interest, and complicates the assessment of the pharmacologically induced responses and is itself of considerable interest. In particular, the reduction in synapse number observed with DOI and LSD following pretreatment with pertussis toxin suggests that G_i/o_ protein-dependent signalling may contribute to the maintenance of synaptic responses in a ligand-specific manner. Although the basis of these effects remains unclear, these findings point to a more complex role of G-protein signalling in functional plasticity and raise questions that merit further exploration.

With respect to immediate-early gene induction, our uncoupling experiments indicate that *c-Fos* and *Egr-2* do not report equivalent downstream signalling states. Whereas *c-fos* showed a comparatively limited sensitivity to G-protein uncoupling, *Egr-2* proved more consistently dependent on both G_q/11_- and G_i/o_ protein-mediated signalling. In this regard, our findings are broadly consistent with the recent study by Xu et al., in which inhibition of G_i/o_ protein signalling attenuated *Egr-2* induction by psychedelics in primary cortical neurons, supporting a role for non-canonical Gi signalling in psychedelic-associated transcriptional responses [45]. However, our data also show substantial sensitivity of *Egr-2* to G_q/11_ protein uncoupling, and therefore favor a more context-dependent interpretation in which both G_i/o_- and G_q/11_ protein-dependent pathways contribute to early gene induction. Particularly noteworthy is the profile observed for DOI, and to a lesser extent LSD, following PTX pretreatment, as these ligands also showed reduced synaptogenesis under the same conditions. Although this association cannot be interpreted as causal, it raises the possibility that *Egr-2* may more faithfully reflect signalling events linked to functionally relevant neuroplastic adaptations than *c-Fos* alone.

A final point concerns the lactate response, which in our previous phosphoproteomic study was associated with a hallucinogen-selective signalling signature and with FOXK2 phosphorylation [26]. In the present work, we extend those observations by showing that lactate accumulation induced by hallucinogenic serotonergic psychedelics is dependent on 5-HT_2A_ receptor expression and is generally suppressed by uncoupling either G_q/11_ or G_i/o_ protein signalling, indicating that both pathways contribute to this metabolic output. The principal exception was DOI, for which PTX enhanced rather than reduced lactate production. This finding contrasts with the model proposed by Xu et al., in which Gi protein-dependent signalling is considered essential for hallucinogenic effects [45], but is more readily reconciled with earlier observations that PTX-sensitive Gi/o protein signalling can, in some contexts, restrain hallucinogen-driven 5-HT2A receptor-dependent responses rather than simply mediate them [46]. One possible explanation is that, in the case of phenethylamine psychedelics such as DOI, Gi/o protein activation may partially limit Gq/11-dependent signalling, such that uncoupling Gi/o permits a greater net response through the pathway linked here to lactate production. This interpretation is also consistent with the greater selectivity of phenethylamines for 5-HT_2_ family receptors [3, 17], in contrast to tryptamines and ergolines, which more readily engage additional serotonergic receptors, including 5-HT_1_ subtypes that preferentially couple to Gi/o proteins. Within this framework, the distinctive behavior of DOI in the lactate assay, together with its selective phosphoproteomic profile in our dataset (Supplementary Table 2)[26], supports the view that ligand-specific allocation of signalling across G_q/11_- and G_i/o_-dependent pathways may be an important determinant of downstream metabolic and hallucinogenic-like responses. Taken together with our previous phosphoproteomic data linking lactate production to compounds active in the mouse head-twitch response assay, these findings further support the view that lactate accumulation may represent a metabolic output associated with hallucinogenic-like signalling in this model.

In conclusion, the present findings support a model in which serotonergic psychedelics engage an integrated signalling network involving 5-HT_2A_ receptors, TrkB-dependent mechanisms and ligand-specific recruitment of G-protein pathways. Within this framework, 5-HT_2A_ receptor expression appears necessary not only for classical early transcriptional and metabolic responses to hallucinogenic ligands, but also for efficient engagement of neurotrophic and neuroplastic programmes that interact functionally with TrkB signalling. At the same time, the differential profiles observed across dendritogenesis, synaptogenesis, immediate-early gene induction and lactate production indicate that these endpoints do not report a single common signalling state, but rather distinct and only partially overlapping components of psychedelic action. The neural stem cell-derived model described here therefore provides a valuable experimental platform with which to dissect the molecular pharmacology of serotonergic psychedelics in a cellular context with atributes closer to the native state, and may prove valuable for distinguishing signalling features associated with hallucinogenic, neuroplastic and metabolic responses to these compounds.

## Acknowledgments

This research was supported by the Spanish Ministry of Science, Innovation and Universities (MCIN) and the State Research Agency (AEI, grant PID2021-125448OB-I00), co-funded by the European Regional Development Fund (FEDER, “A way of making Europe”) (JFLG). NIH R01MH084894 (JGM) and NIH T32DA007027 (JLM).

## Conflict of interest

Authors declare that they have no conflict of interest.

---

**Supplementary Fig.1.**
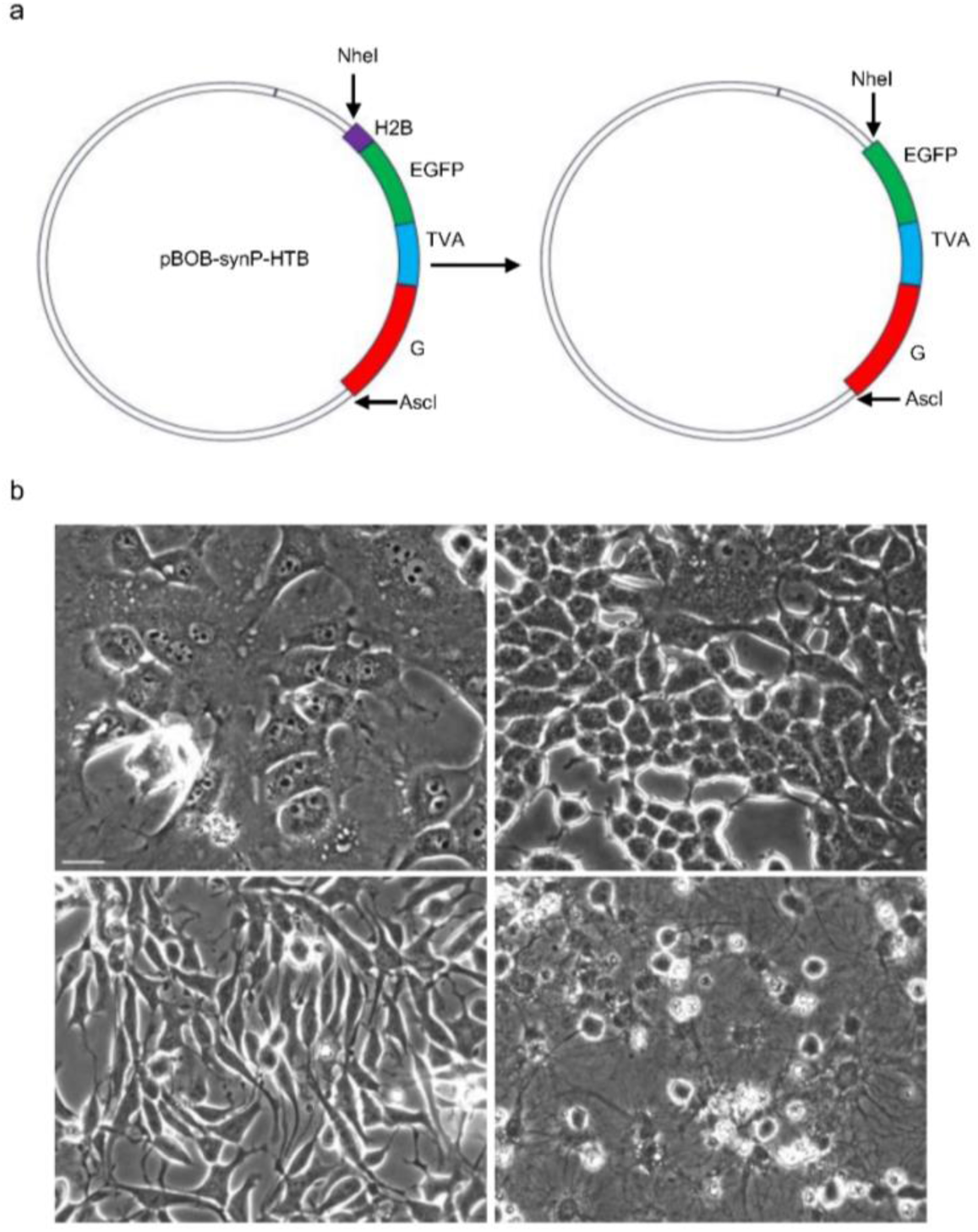
Strategy for the removal of the H2B sequence from pBOB-synP-HTB. **a.** The original plasmid pBOB-synP-HTB (left) contained the H2B-EGFP-TVA-G cassette. To eliminate the H2B coding sequence, the plasmid was digested with NheI and AscI. In parallel, the fragment containing EGFP-TVA-G was PCR-amplified using primers introducing NheI and AscI restriction sites at the 5’ and 3’ ends, respectively. The amplified fragment was ligated into the digested backbone, yielding the modified plasmid in which the H2B sequence was replaced by the EGFP-TVA-G cassette (right). **b.** Upper panels: Neural stem cells (NSCs) expressing the original pBOB-synP-HTB plasmid before (left) and after five days of differentiation (right). Lower panels: NSCs expressing the modified plasmid lacking the H2B sequence before (left) and after five days of differentiation (right). Scale Bar: 20 µm.

**Supplementary Fig. 2.**
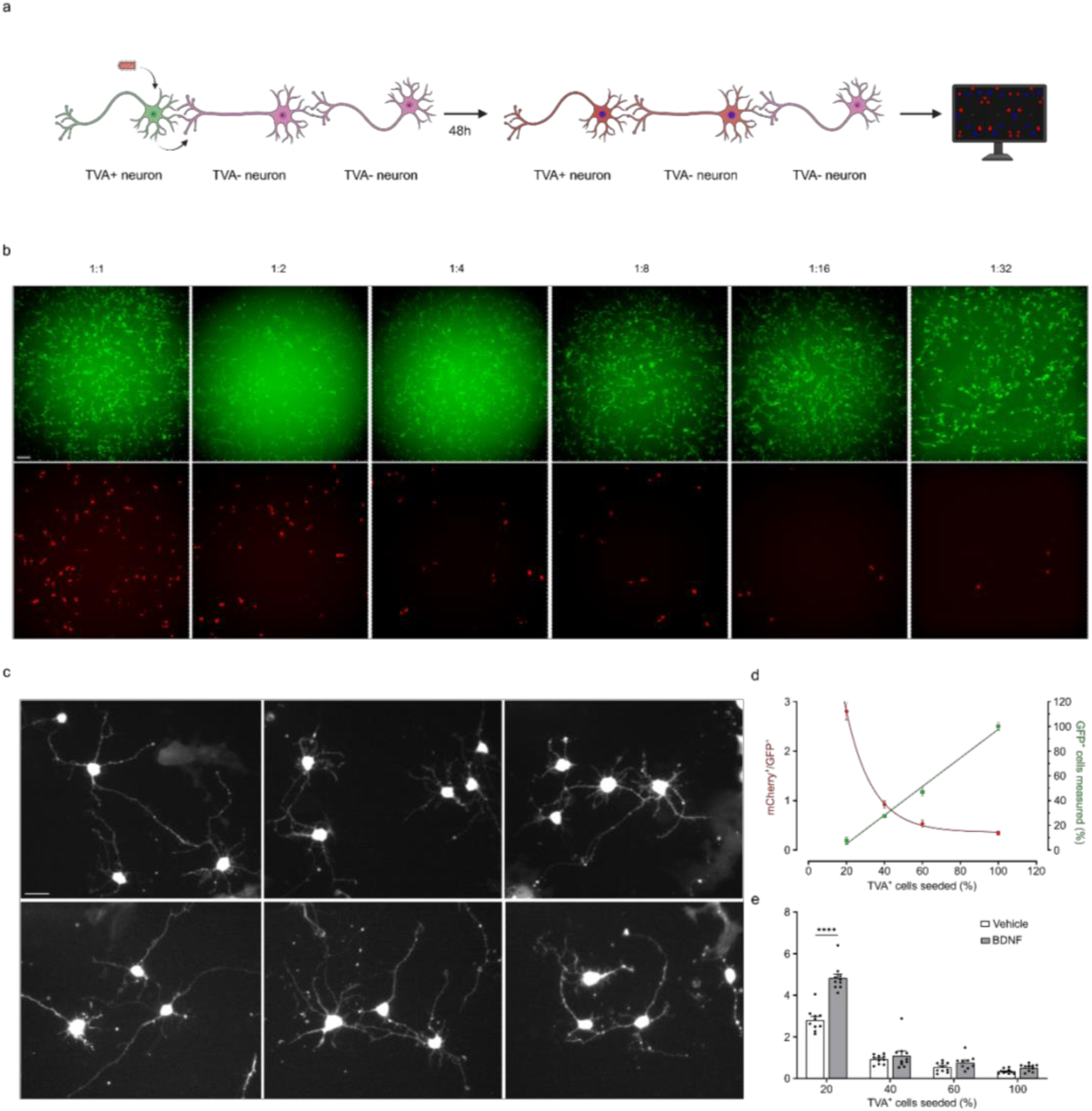
Development of an *in vitro* methodology to evaluate synaptogenesis using rabies virus neuronal tracing technology. **a.** Schematic representation of the assay used to quantify synaptic connectivity by rabies virus neuronal tracing. Rabies virus particles (in red), encoding the mCherry fluorescent reporter and pseudotyped with the EnvA envelope glycoprotein, selectively infect NSC TVA derived neurons (in green) that express the TVA receptor and EGFP. The virus is then transmitted retrogradely to presynaptic NSC shRNA TrkB or NSC shRNA 5-HT_2A_ derived neurons (in pink). Synaptic inputs are quantified by counting mCherry-positive (red) cells using image analysis. **b.** Undifferentiated neural stem cells (NSCs) expressing TVA receptors, EGFP, and the rabies glycoprotein were selected by flow cytometry based on EGFP fluorescence. Cells were then infected with different dilutions of EnvA-pseudotyped rabies virus expressing mCherry, which appears as red fluorescence in the lower panels. Images were acquired at 10x magnification. **c.** Representative microscopic fields of neurons derived from NSC TVA differentiation, exhibiting mCherry fluorescence 48 hours post-rabies virus infection. Scale bar: 20 µm. **d.** Evaluation of the proportion of NSC TVA cells seeded in mixture with wild type NSCs to achieve an optimal ratio of total mCherry expressing neurons relative to EGFP expressing neurons. Left axis: ratio of mCherry+ to EGFP+ cells. Right axis: percentage of EGFP+ cells. **e.** Evaluation of the proportion of NSC TVA cells seeded in mixture with wild type NSCs to identify an optimal window for quantifying the effects of neuroplastogenic treatments on mCherry+ neuron numbers. Cells were treated with BDNF (50 ng/mL) for 24 h, and synaptogenesis was assessed according to the established protocol (see Methods). Data were analyzed by two-way ANOVA, revealing significant effects of cell proportion (F(3,71) = 280.1, P < 0.0001), treatment (F(1,71) = 48.03, P < 0.0001), and their interaction (F(3,71) = 23.43, P < 0.0001). ****p < 0.0001.

**Supplementary Fig. 3.**
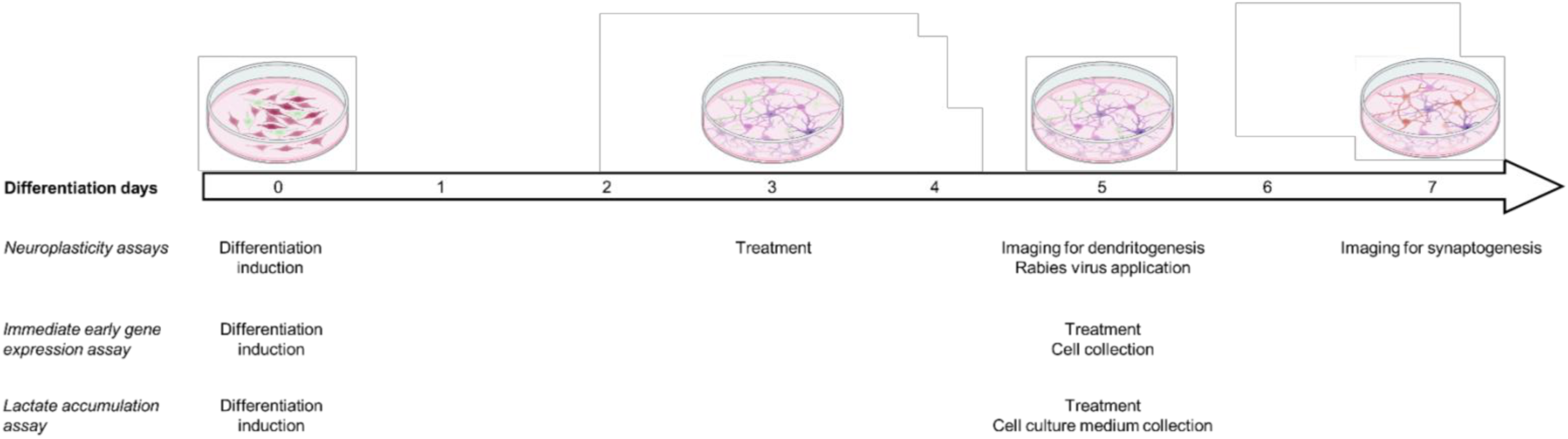
Experimental timelines. Schematic representation of the experimental timeline during the differentiation stage of the cultures used for neuroplasticity, immediate early gene expression, and lactate accumulation experiments.

**Supplementary Fig. 4.**
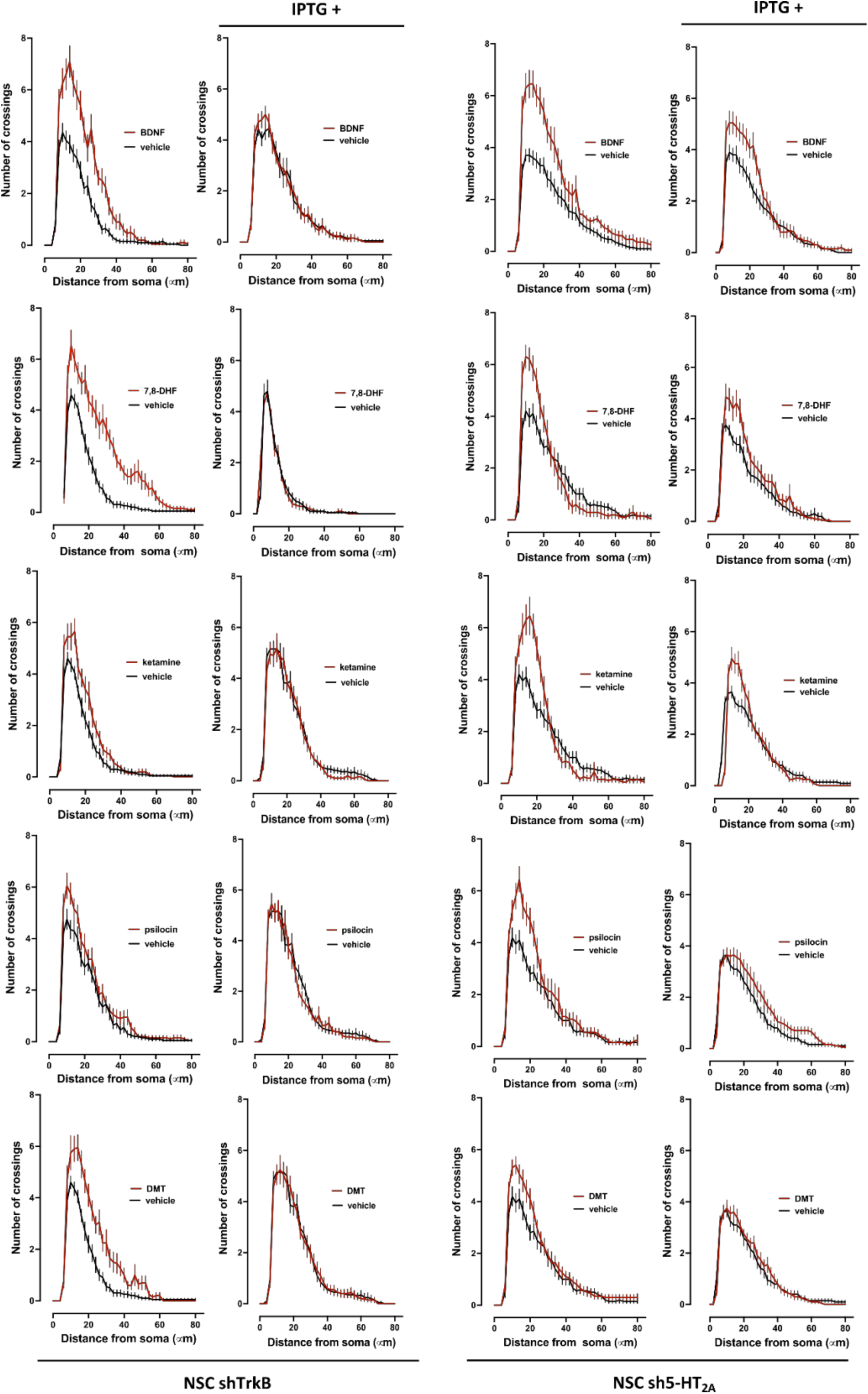
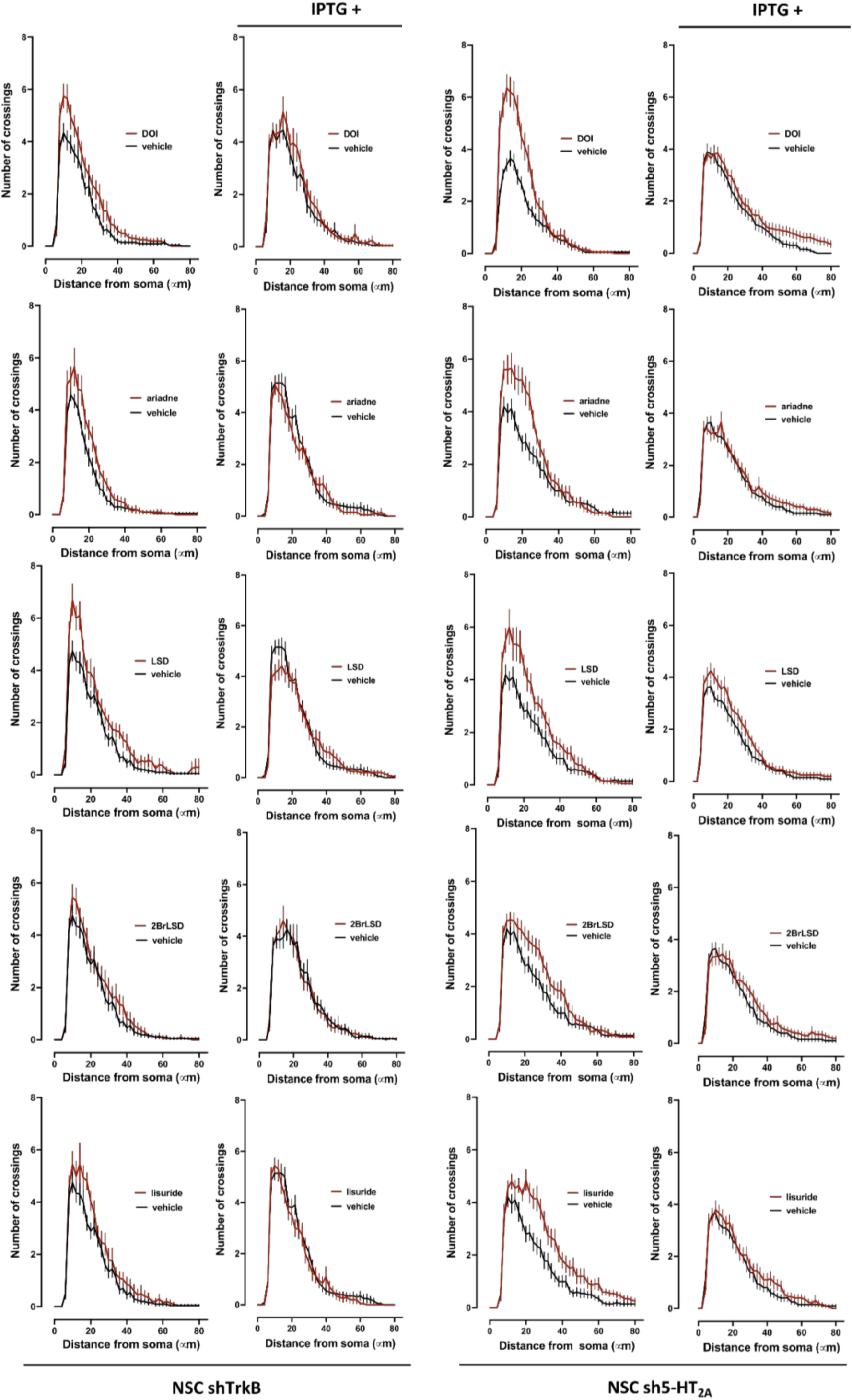
Effect of drug treatments and receptor silencing on dendritic arborization complexity assessed by Sholl analysis. Graphs illustrate a representative neuroplasticity experiment showing Sholl curves from neurons derived from NSC shTrkB or NSC sh5-HT_2A_ cell lines treated with the indicated compounds, compared with their respective vehicle controls. Columns designated as IPTG+ correspond to cells pretreated with IPTG to silence the expression of either TrkB or 5-HT_2A_ receptors.

**Supplementary Figure 5.**
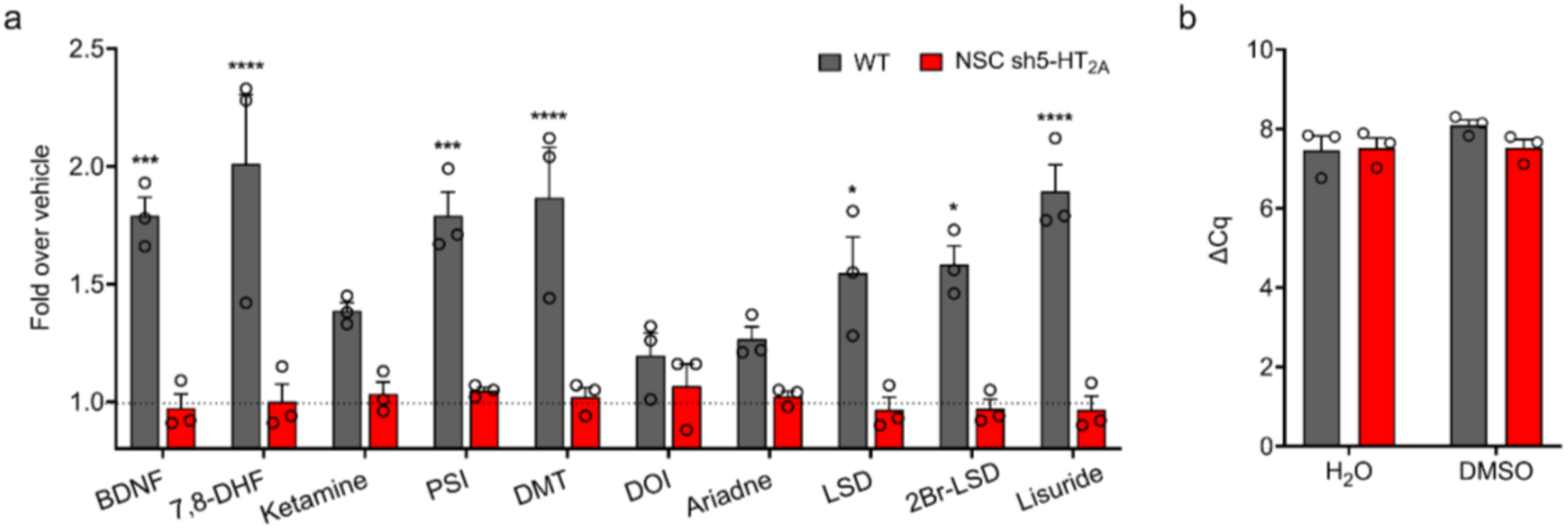
Effect of drug treatments and 5-HT_2A_ silencing on *Bdnf* mRNA transcription. Wild type NSCs (grey bars) and 5-HT_2A_-silenced NSCs (NSC shRNA 5-HT_2A_, red bars) were differentiated into neurons and glia and treated with the indicated drugs to **a:** Evaluate the transcription of *Bdnf* gene represented as changes relative to basal levels. Two-way ANOVA revealed significant main effects of drug treatments (F(10,40) = 4.715, p = 0.0001) and cell phenotype (F(1,40) = 166.8, p < 0.0001). A significant interaction was also observed (F(10,40) = 5.436, p < 0.0001). Post hoc comparisons were performed using Tukey’s multiple comparison test. *p < 0.05, **p < 0.01, ***p < 0.001, ****p < 0.0001. **b:** Quantification by RT-qPCR of the *Bdnf* gene transcripts when treating wild type NSCs (grey bars) and 5-HT_2A_-silenced NSCs (NSC sh5-HT_2A_, red bars) with vehicles. No significant differences were observed (H20: p = 0.8895; DMSO: p = 0.1507).

**Supplementary Fig. 6.**
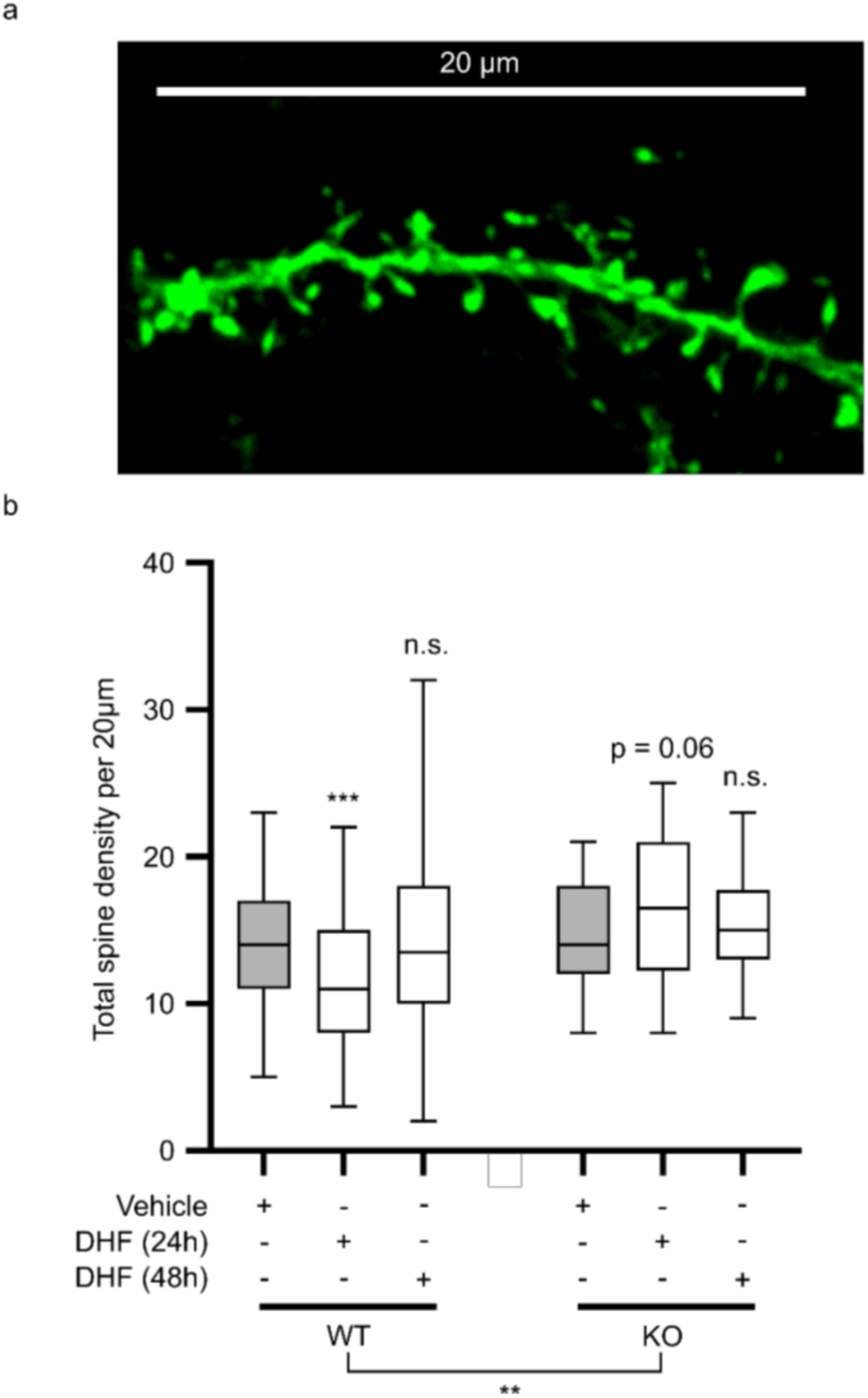
Effect of 7,8-DHF treatment in *5-HT_2A_R-KO* mice and wild-type controls. Effect of chronic 7,8-DHF administration on dendritic spine density in the frontal cortex of *5-HT_2A_R-KO* mice and wild-type controls. **a.** Representative three-dimensional reconstructions of AAV-injected frontal cortex dendritic segments. Scale bar represents 20 µm. **b.** Dendritic spine density in wild-type (n = 5-75 segments from 2-4 mice) and *5-HT_2A_R-KO* (n = 10 segments from 2-3 mice) animals. Statistical analysis was performed using two-way ANOVA (genotype F(1,375) = 9.060, p = 0.0028; DHF F(2,375) = 0.314, p = 0.7305; interaction F(2,375) = 5.735, p = 0.004) (**p < 0.01), or one-way ANOVA (WT: F(2,308) = 11.06 p < 0.0001; *KO*: F(2,67) = 1.781 p = 0.1764) followed by Bonferroni’s multiple comparison test (***p < 0.001, n.s., not significant).

**Supplementary Fig.7.**
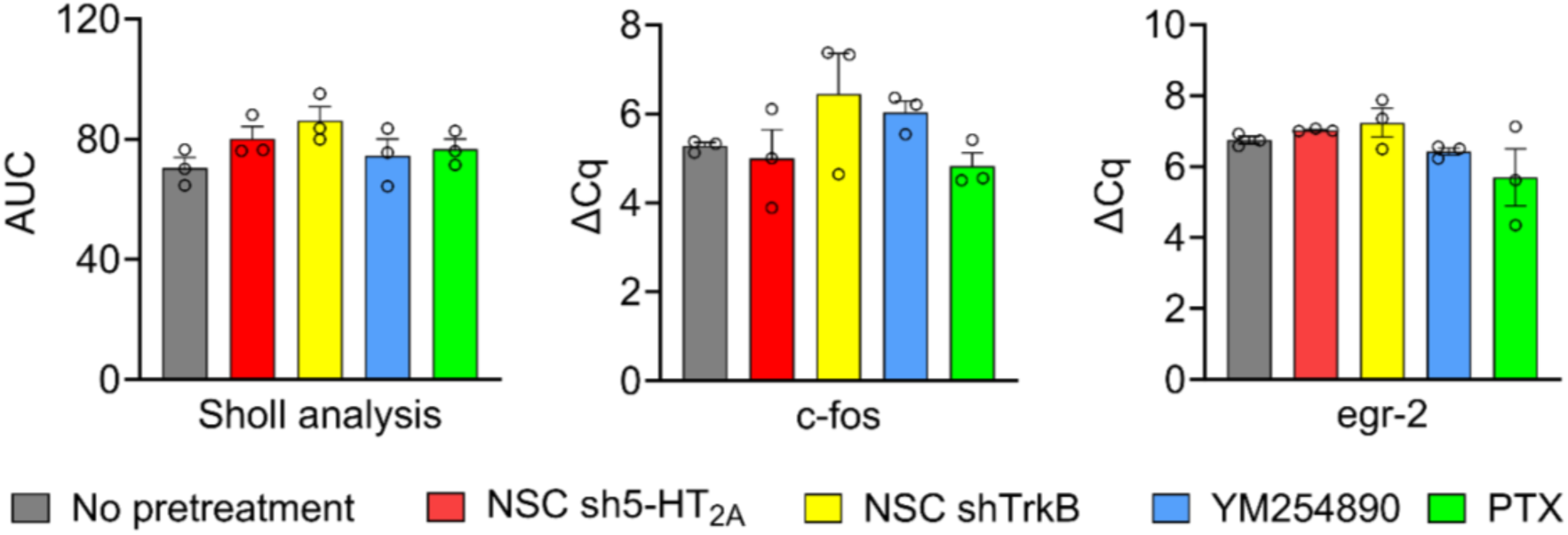
Silencing and decoupling G proteins does not affect basal dendritogenesis and immediate early gene expression. Measurement of the effect of decoupling G proteins, and silencing *Ntrk2* and *Htr2a* genes on: **a.** AUC of the neurites measured through Sholl analysis when treating cells with vehicle. One-way ANOVA, no significant effects were detected for the interaction (F(4,10) = 1.983, p = 0.1732). **b.** Quantification by RT-qPCR of *c-fos* expression when treating with vehicles. One-way ANOVA, no significant effects were detected for the interaction (F(4,10) = 1.754, p = 0.2147). **c**. Quantification by RT-qPCR of *egr-2* expression when treating with vehicles. One-way ANOVA, no significant effects were detected for the interaction (F(4,10) = 2.186, p = 0.1440).

**Supplementary table 1.** Peptides differentially phosphorylated when treating with psilocin. The main theme of the functions of these proteins is cytoskeletal and microtubule regulation. FDR = 0.999958303.

| Peptide | Phosphorylations | Mean difference | p value | Z score |
| --- | --- | --- | --- | --- |
| Fam169a | (S630); (S631) | -2.3423858 | 0.11315347 | -4.6582645511745 |
| Usp9x | (S588) | -2.41093023 | 0.14810003 | -4.7923170926679 |
| Rras2 | (S186) | 2.34870436 | 0.00243058 | 4.51611299238523 |
| Nes | (S1444) | 2.97031606 | 0.00199617 | 5.73180066761452 |
| Map2 | (T1445) | -2.83264085 | 0.08740626 | -5.6170576661681 |
| Hdgf | (S128); (S132); (S133) | 2.6404283 | 0.0264845 | 5.08663829191033 |
| Numa1 | (S2059) | -2.46464204 | 0.01737692 | -4.8973614077365 |
| Bcas1 | (S347) | -2.6928905 | 0.02483944 | -5.3437475459567 |
| Mast2 | (S1178); (K1175) | 2.72976459 | 0.02375465 | 5.26135350595555 |
| Tns1 | (R1532); (S1535) | 2.36715086 | 0.03577025 | 4.55218887422095 |
| Map1b | (R1660); (S1686); (S1671) | -4.05343382 | 0.00135835 | -8.0045657367117 |
| Foxk1 | (S87); (R85) | -2.83027966 | 0.0017735 | -5.6124398904469 |
| Fez2 | (S200) | 2.5294581 | 0.0304026 | 4.86961356576167 |
| Mapre2 | (S222) | -2.53224934 | 0.21068761 | -5.0295812022094 |

**Supplementary table 2.**
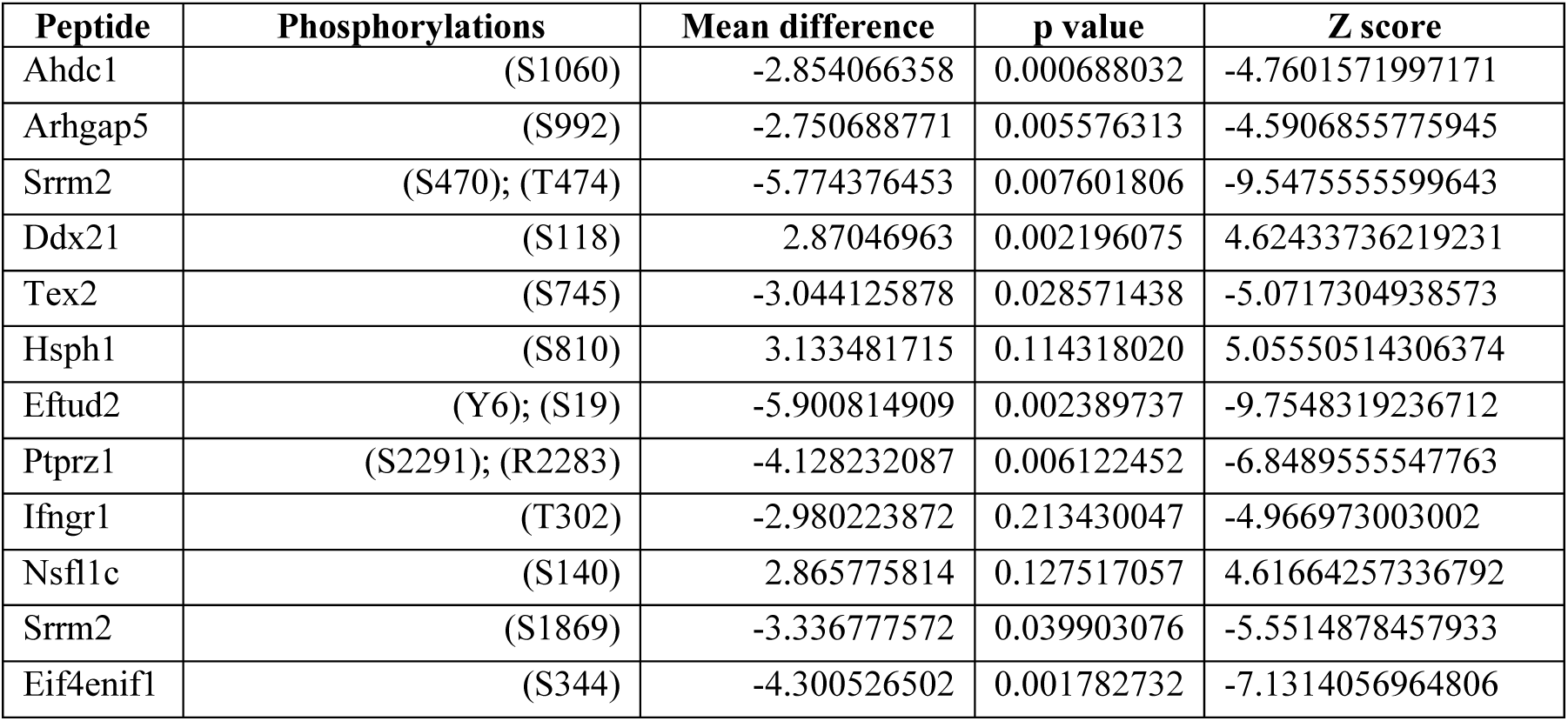
Peptides differentially phosphorylated when treating with DOI. The main theme of the functions of these proteins is transcriptional regulation through nucleolar activity, and RNA processing and trafficking. FDR = 0.934992069.

## Notes

### Competing Interest Statement

The authors have declared no competing interest.

